# Genetic and geometric heredity interact to drive polarized flow in the *Drosophila* embryo

**DOI:** 10.1101/2022.07.13.499934

**Authors:** Emily Gehrels, Bandan Chakrabortty, Matthias Merkel, Thomas Lecuit

## Abstract

Tissue flow during morphogenesis is commonly driven by local constriction of cell cortices, which is caused by activation of actomyosin contractility. This can lead to long-range flows due to tissue viscosity. However, in the absence of cell-intrinsic polarized forces or polarity in forces external to the tissue, these flows must be symmetric and centered around the region of contraction. Polarized tissue flows have been previously demonstrated to arise from the coupling of such contractile flows to points of increased friction or adhesion to external structures. However, we show with experiments and modeling that the onset of polarized tissue flow in early *Drosophila* morphogenesis occurs independent of adhesion and is instead driven by a geometric coupling of apical actomyosin contractility to tissue curvature. Particularly, the onset of polarized flow is driven by a mismatch between the position of apical myosin activation and the position of peak curvature at the posterior pole of the embryo. Our work demonstrates how genetic and geometric information inherited from the mother interact to create polarized flow during embryo morphogenesis.

## Introduction

Morphogenesis is the process by which organisms develop from a simple fertilized egg to an adult with complex form and function. This process depends keenly on the dynamics of the underlying biological tissues, which itself arises from cellular attributes such as cell-cell adhesion, cortical tension, osmotic pressure, elasticity, and viscosity, all of which can be passively and actively controlled in space and time^1,2^. Tissue-scale effects can arise as a result of local changes to these cellular attributes. For example, tissue invagination emerges from apical constriction of well-defined groups of cells^3^ such as in the mesoderm or endoderm of *C. elegans*^4,5^, *Drosophila*^6^, sea urchin^7^, or ascidians^8^ (also reviewed in Ref. 9).

Tissue flows are ubiquitous in animal development, especially during embryogenesis and organogenesis. Tissue flow may be symmetric or polarized (asymmetric and vectorial) depending on the pattern and polarity of internal active stresses and on the existence of external polarized active stress acting on the tissue^2^. When no external polarized force exists, flow is intrinsically symmetric. Cells may converge towards a group of constricting cells such as on either side of the *Drosophila* mesoderm^10^. Alternatively, anisotropic forces driving cell division can give rise to local divergent flow due to bipolar cell displacement^11,12^ or tissue extension^13^. These two types of symmetric flows may coexist, perpendicular to each other, during the process of so-called convergent extension. During this process, cell intercalation causes local divergent flow and perpendicular convergence due to junction shrinkage along one axis, and junction extension along the perpendicular axis^14–21^. In contrast, the emergence of polarized flow depends upon the presence of an external acting force. For instance, contraction of the hinge at the proximal end of the developing *Drosophila* pupal wing and anchoring of the wing at its distal tip drive polarized tissue flow and global extension of the wing^12,22,23^. Similarly, a supracellular tensile ring in the posterior of chick epiblast drives non-local polarized rotational flows in the whole embryonic field during primitive streak formation^24^.

During early *Drosophila* morphogenesis, a single layer of epithelial cells is formed by the simultaneous cellularization of the 6000 nuclei composing the syncytial blastoderm^25^. Subsequently, at the onset of gastrulation, this initially static tissue begins to flow and thereby initiates the process of axis elongation along the antero-posterior axis^14,15^. Most notably, the tissue in the posterior of the embryo, called the endoderm, undergoes a polarized flow towards the dorsal-anterior side of the embryo^16,26^ (**Supplementary Video 1, Fig. 1a**). This sharp onset of polarized flow makes the *Drosophila* embryo a powerful system to study the physical mechanisms that drive tissue flow.

**Figure 1:**
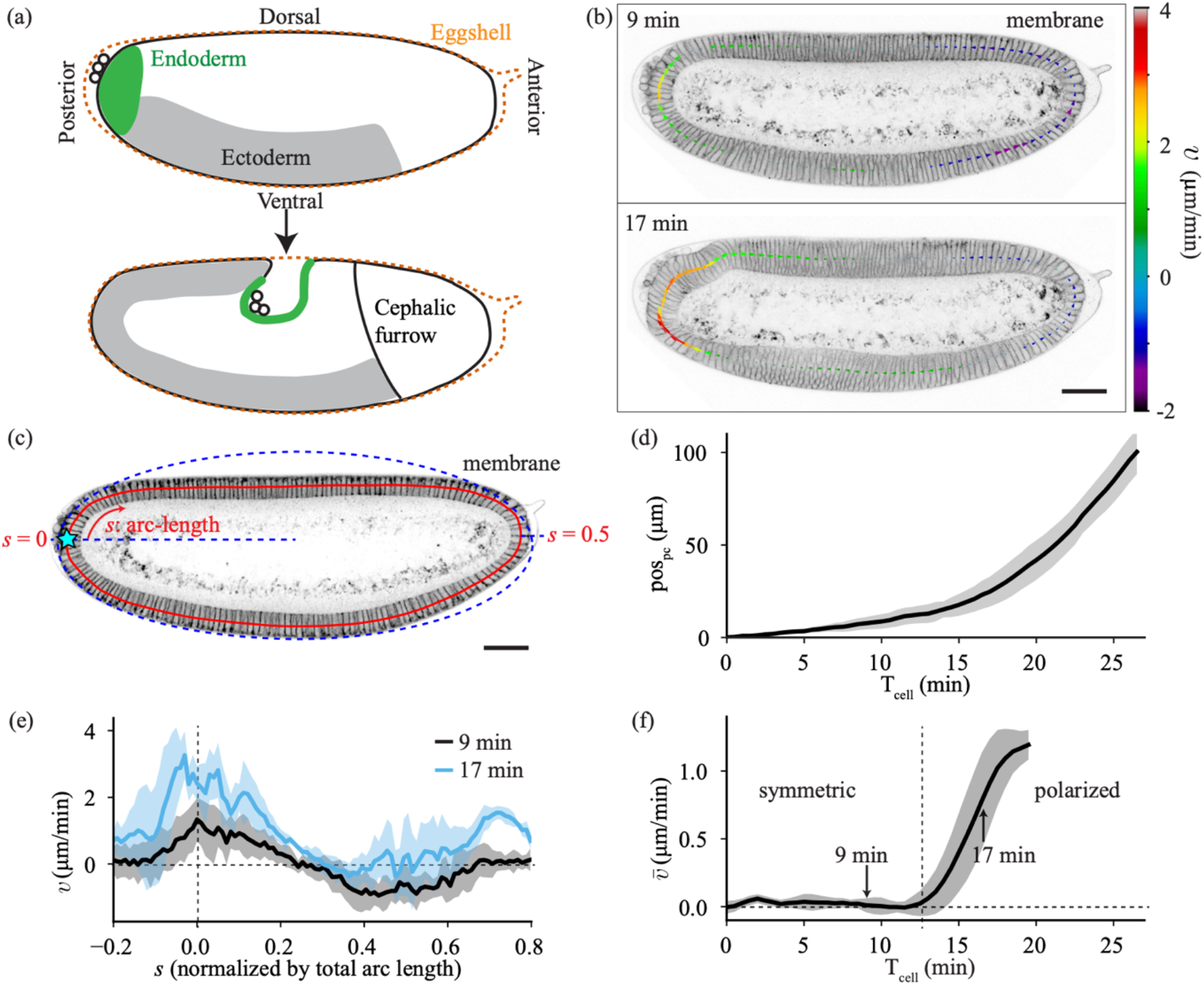
Quantification of tissue flow during early *Drosophila* morphogenesis. **(a)** Cartoons of *Drosophila* embryo (*top*) at an early stage, during the process of cellularization and (*bottom*) approximately 30 minutes later. **(b)** Sagittal plane of an embryo imaged with membrane marker GAP43::mScarlet at 9 minutes and 17 minutes after the cellularization front passes the nuclei in the dorsal posterior (T_cell_ =0). The arrows show the tangential velocity of the tissue along the midline, extracted using Particle Image Velocimetry (PIV). **(c)** Two-photon images of a cut through the sagittal plane of the embryo imaged with the membrane marker GAP43::mScarlet. The red line denotes the midline of the epithelium. The arc-length variable *s* increases in clockwise direction and is normalized such that *s* = 1 corresponds to the total length of the embryo midline. The blue dashed line represents a fit of an ellipse to the apical surface of the epithelium, which is used to determine the location of *s* = 0 at the posterior pole (see Methods). **(d)** Quantification of the position of the pole cells (pos_pc_, see Methods) as a function of time after the cellularization front passes the nuclei in the dorsal posterior (T_cell_). Average performed over 6 embryos. **(e)** Spatial profile of tangential tissue velocity along the midline at the same times as in **c**. Average performed over 5 embryos. **(f)** Spatial average of the tangential velocity 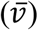 as a function of time. Vertical dashed line marks the onset of polarized flow. Average performed over 5 embryos. All scale bars are 50 μm. Error bars represent the standard deviation.

It is so far unclear what creates this early polarized flow in the *Drosophila* embryo. One possibility is that it might be coupled to and dependent on other processes during early *Drosophila* morphogenesis, such as mesoderm invagination on the ventral side^10^, germband extension along the lateral side^16^, or the cephalic furrow in the anterior^27^. Indeed, existing studies have shown that most of the embryonic tissue flow can be accurately predicted from the pattern of non-muscle Myosin-II (hereafter myosin) localization and polarization^28^. However, earlier studies revealed that an additional force arising near the posterior pole must contribute to driving tissue flows, as evidenced by gradients of cell shape^16,26^ and mechanical stresses^16^, as well as genetic and mechanical perturbations^16,29^. For instance, blocking endoderm invagination (as in a *Torso* mutant) entirely blocks whole-embryo elongation^16^. Moreover, the early polarized flow of the endoderm is independent of the above-mentioned processes, as demonstrated by embryos with simultaneously blocked mesoderm invagination (*twist, snail* ^[30]^), germband extension, and cephalic furrow formation (*eve* ^[31]^; **Supplementary Video 2, Extended Data Fig. 1a, b**). Hence, the question remains as to what causes the first polarized tissue flow of the *Drosophila* epithelium and initiation of axis elongation.

### *Drosophila* morphogenesis begins with symmetric tissue flow that becomes polarized

To understand what drives tissue flow in early *Drosophila* morphogenesis, we first quantified the flow to determine how it evolves in time. To do so, we performed live imaging of *Drosophila* embryos using a two-photon microscope to capture the sagittal plane that cuts through the center of the embryo (**Fig. 1b, c, Methods**). We tracked the position of the pole cells (pos_pc_) to quantify the motion of the posterior tissue (**Supplementary Video 3, Methods**), which revealed that there is an initially slow flow of the posterior, which speeds up over time (**Fig. 1d**).

We also quantified the spatial profile of the flow in the epithelium by performing particle image velocimetry (PIV) on subsequent frames and extracting the velocity of the flow tangential to the midline of the epithelium (*v*; **Fig. 1b, c, Methods**). At each time, the tangential velocity (**Fig. 1e**) can be spatially averaged to give a single value that describes the global polarity of the flow (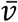; **Fig. 1f**). If there is as much clockwise (positive) flow as counterclockwise (negative) flow, the average is zero, and the flow is considered symmetric (as at T_cell_ = 9 min). If there is more clockwise flow than counterclockwise flow, the average will be positive, indicating that the flow is polarized (as at T_cell_ = 17 min). Averaging the velocity revealed that there are two distinct phases of flow: a symmetric flow for approximately the first twelve minutes after the cellularization front passes the nuclei (T_cell_ = 0, **Methods**), followed by the onset of polarized flow with average velocity that increases in time (**Fig. 1f**).

When we performed the same quantifications on embryos mutant for *eve, twist*, and *snail* (hereafter *ets*), which have blocked mesoderm invagination, germband extension, and cephalic furrow formation (**Supplementary Video 2, Extended Data Fig. 1a, b**), we still saw this transition to polarized flow (**Extended Data Fig. 1c, d**). The timing of the transition was even slightly earlier in *ets* embryos than in wildtype, likely due to the lack of ventral pulling from the mesoderm invagination, as evidenced by the increased flow in the ventral-anterior region of the embryo (−0.2 < *s* < 0; **Extended Data Fig. 1e, f**). This clearly demonstrates that the polarized flow does not depend on myosin polarization in the germband, or on geometric constraints imposed by the cephalic furrow as previously predicted^27,28^. This lead us to two main questions: What physical mechanisms drive tissue flow in the early *Drosophila* embryo? And what mechanism is involved in the transition from symmetric to polarized flow?

### Symmetric and polarized tissue flows arise from basal and apical myosin respectively

Myosin is known to be a common tissue-intrinsic driver of flow in many biological tissues^1,32^. We therefore imaged the myosin distribution in the embryo as a function of time to determine its impact on tissue flow. In wildtype embryos (**Fig. 2a** top row), there are two distinct populations of myosin: one on the apical side of the cells, and the other on the basal side.

**Figure 2:**
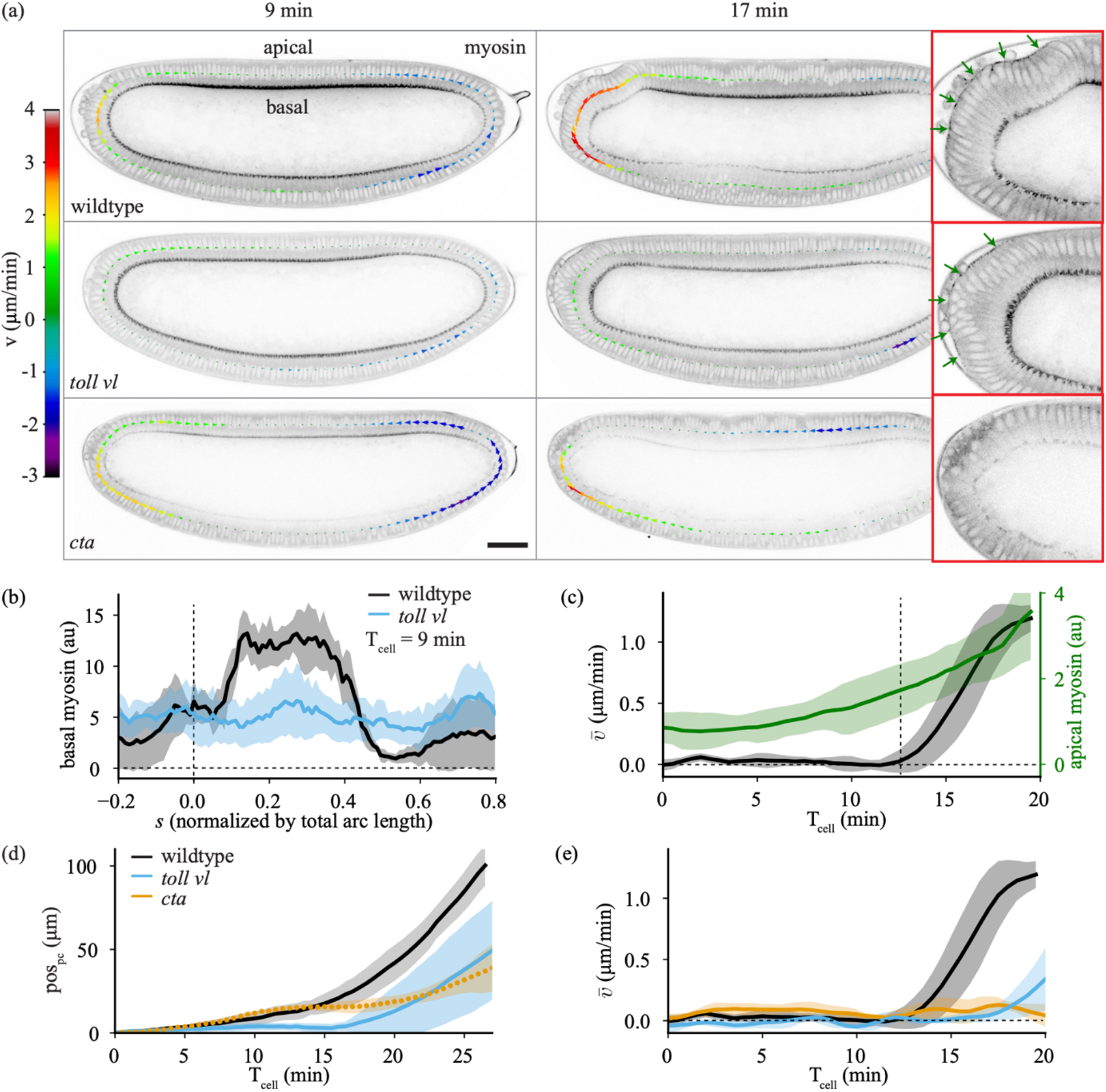
Impact of apical and basal myosin on tissue flow. **(a)** Sagittal plane of an embryo imaged with Myosin-II (myosin) marker *spaghetti squash* (*sqh*)::GFP at 9 minutes and 17 minutes after T_cell_ = 0 for (*top*) wildtype (*middle*) *toll vl* mutant and (*bottom*) *cta* mutant embryos. The arrows show the tangential velocity of the tissue along the midline. The panels on the right show a zoomed view of the posterior at T_cell_ = 17 min with apical myosin indicated with green arrows. (**b**) Spatial profile of basal myosin intensity at T_cell_ = 9 min for wildtype (black) and *toll vl* mutant (blue) embryos. Average performed over 5 wildtype and 6 *toll vl* mutant embryos. (**c**) Average of apical myosin over the posterior of the embryo (−0.1 < *s* < 0.15; green) and spatial average of the tangential velocity as a function of time (black) for 5 wildtype embryos. Vertical dashed line marks the onset of polarized flow. (**d**) Pole cell position (pos_pc_) as a function of time for wildtype, *toll vl*, and *cta* embryos. Average performed over 6 wildtype, 7 *toll vl*, and 7 *cta* embryos. (**e**) Spatial average of the tangential velocity as a function of time. Average performed over 5 wildtype, 6 *toll vl*, and 5 *cta* embryos. All scale bars are 50 μm. Error bars represent the standard deviation.

During the symmetric phase of flow, there is no apical myosin activation, but there is a strong accumulation of basal myosin (**Fig. 2a** top left), associated with the process of cellularization^25^. Cellularization completes first on the ventral side of the embryo, causing more basal myosin dorsally than ventrally (**Fig. 2b**), which coincides with the center of the domain of symmetric tissue flow (**Extended Data Fig. 2a**). This lead us to the hypothesis that non-uniformity of basal myosin could drive the symmetric phase of tissue flow, as previously proposed^28^.

To test this hypothesis, we used a mutation that removes the dorsal-ventral specification of the embryo, and leads the entire embryo to act as ventrolateral tissue^33^ (*toll*^rm9^/*toll*^rm10^ hereafter *toll vl*; **Fig. 2a** middle row). This makes the embryo rotationally symmetric about its anterior-posterior axis (**Extended Data Fig. 2b**), which in turn removes the dorsal-ventral difference in basal myosin (**Fig. 2b, Supplementary Video 5**). In *toll vl* embryos, the posterior tissue does not always flow dorsally, but instead is significantly more likely than wildtype to flow towards the ventral or lateral directions (**Extended Data Fig. 2c, Supplementary Video 4**). To quantify the flow in the imaging plane, we analyzed only embryos that flow dorsally. Tracking the pole cells in these mutants revealed that the early, symmetric flow is almost completely halted (**Fig. 2d**), confirming that the symmetric flow observed is driven by the increased activation of basal myosin on the dorsal side of the embryo.

We next considered what drives the polarized flow that occurs following the symmetric phase of flow. Around the time where the transition occurs, the levels of basal myosin begin to decrease and there is a localized accumulation of apical myosin in the dorsal posterior (**Fig. 2a top right, c**). To test whether the polarized flow requires this posterior pool of apical myosin, we strongly downregulated all apical myosin in the embryo using a mutation in the gene encoding for the G protein Gɑ_12/13_ (known as Concertina (Cta) in *Drosophila*), which is required specifically for medial apical myosin recruitment^34,35^ (**Fig. 2a** bottom row, **Supplementary Video 5**). Quantifying the flow by pole cell tracking (**Fig. 2d**) and by averaging epithelial velocity (**Fig. 2e**) showed that the symmetric flow is similar to wildtype, but that the polarized flow is strongly suppressed.

Based on these observations, we concluded that the symmetric phase of flow requires the nonuniformity of basal myosin and the polarized flow requires apical myosin. Since polarized tissue flow is normal in *ets* mutants in which mesoderm invagination and germband extension are blocked, we concluded that apical myosin is required strictly in the posterior region of the embryo and not in other adjacent tissues. It is known that the main effect of myosin accumulation is to drive contraction of the acto-myosin network inside of cells^1,32^. We therefore decided to investigate whether the observed flow dynamics arise solely due to tissue contraction driven by these two different pools of myosin.

### Myosin-driven tension-based model explains the symmetric flow

To gain mechanistic insight into the process driving the flow, we decided to test our hypotheses using a combination of experiments and modeling. Our aim was to understand the embryo flow dynamics on the tissue scale, we therefore chose to use a model that considers collective cellular behavior rather than individual cellular processes. Because the strain rates measured during this phase of development do not surpass approximately 1/50 min^-1^ (**Extended Data Fig. 3a, b**), we neglected elastic stresses, which are expected to relax much faster, on the time scale of approximately 2-5 min ^[36]^. Finally, we focused on the flow within a sagittal section of the embryo, which we described by a continuous, thin 1D membrane. These simplifications allowed us to compare our sagittal observations of embryos in two-photon microscopy to a model of a 1D thin active fluid^37,38^ (**Supplementary Information**), which takes into account tissue viscosity, friction with the vitelline membrane (part of the eggshell that surrounds the embryo), and myosin-driven spatially-dependent active tension (equation (1) in **Fig. 3a**).

**Figure 3:**
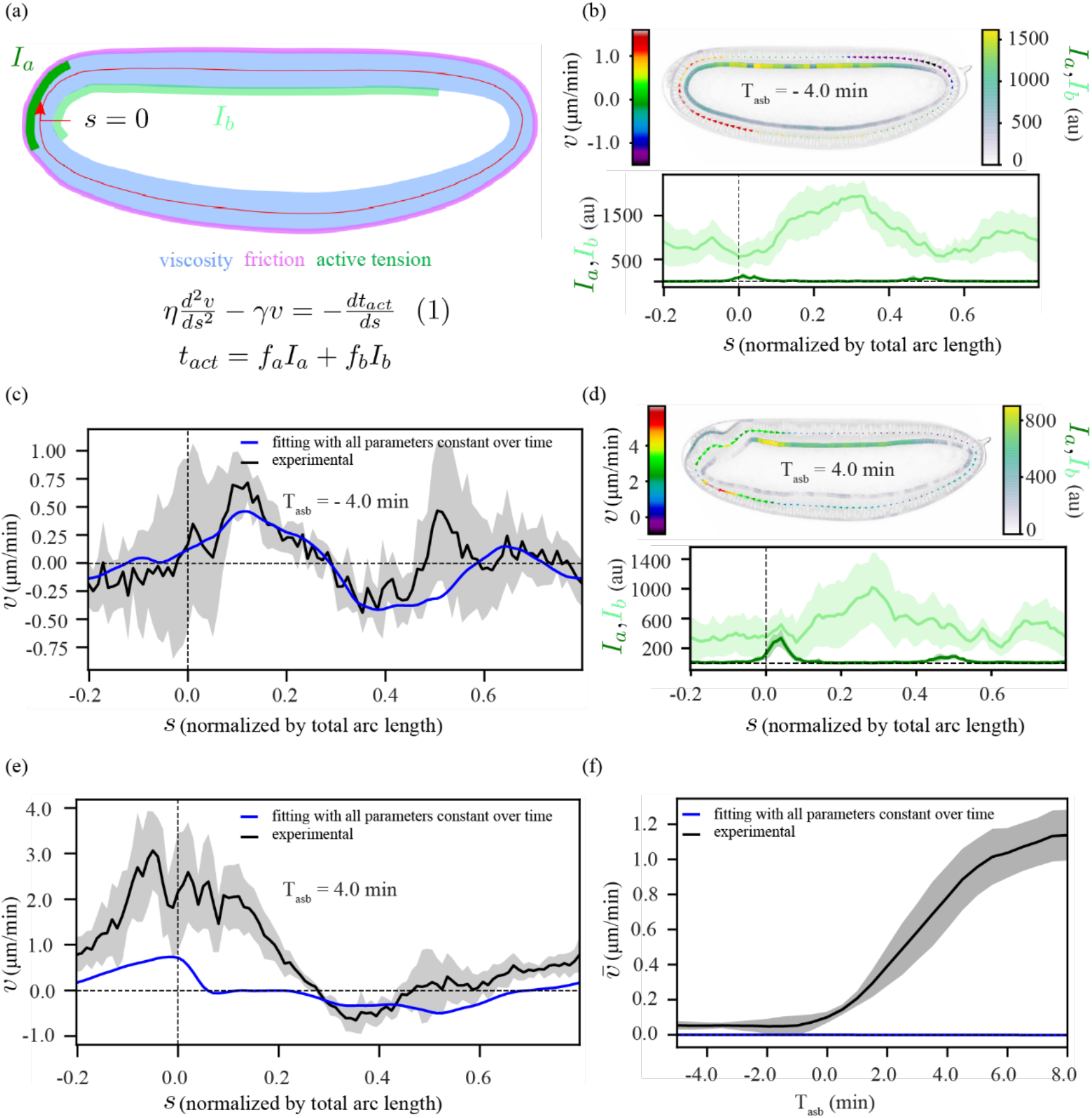
A model based on an active tension mechanism explains the symmetric flow. **(a)** Schematic representation of our modeling framework, equation (1), in which the epithelium is considered as a viscous fluid (*η*, blue) with homogeneous friction (*γ*, pink) with the surroundings, along with domains of apical myosin (*I*_*a*_, dark green) and basal myosin (*I*_*b*_, light green) that act as contractile elements. The tissue is approximated by a 1D continuous membrane, positioned along the midline of the epithelium (red line). At any given position, *s*, the tangential component of the velocity (*v*) fulfills equation (1). **(b, d)** (*top*): a representative time frame from the symmetric phase with a heatmap of *v, I*_*a*_ and *I*_*b*_, (*bottom*): the corresponding spatial profiles of *I*_*a*_ and *I*_*b*_, averaged over 6 *ets* embryos. (**b)** at T_asb_ = -4 min (**d)** at T_asb_ = 4 min. **(c**,**e)** Fit (blue) of equation (1) to the experimentally measured *v* (black). We perform a simultaneous fit to the data for all time points between T_asb_ = -5 min and T_asb_ = 8 min, where we set all parameter values, *l*_*H*_, *r*_*a*_ *= f*_*a*_*/η*, and *r*_*b*_ *= f*_*b*_*/η* constant over time (see Supplementary Information). **(c)** T_asb_ = -4 min, **(e)** T_asb_ = 4 min. **(f)** Temporal profile of spatially averaged velocity 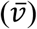 in experiment (black) and spatial average from our simultaneous fit (blue). For further details see **Extended Data Fig. 4** and Supplementary Information. In all panels, the shaded regions indicate the standard deviation, computed over 6 embryos.

This type of model is effective for making quantitative comparisons as well as qualitative predictions. To quantitatively compare our model to the data, we solved equation (1) for velocity, using the measured apical and basal myosin intensities (**Fig. 3b**) as input to the equation. By fitting to the measured velocity field, we extracted the values of three relevant physical parameters (**Supplementary Information**). The first parameter is a hydrodynamic length scale 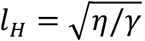, which reflects the ratio between viscosity, *η*, and friction with the surroundings, *γ*. The other two parameters are conversion factors that relate myosin intensity to active tension (*f*_*a*_ for apical myosin and *f*_*b*_ for basal myosin), which are divided by viscosity *η*. The resulting ratios, *r*_*a*_ and *r*_*b*_, reflect tissue contraction rates for apical and basal myosin respectively. To build qualitative intuition, we also performed simulations of a simplified model without basal myosin on an elliptic representation of the embryo (hereafter elliptic model), using values of the relevant physical parameters from our fitting (**Supplementary Information**).

We then used this model to test our hypothesis that the non-uniform contraction of basal myosin is what drives the observed symmetric flow. To simplify our modeling studies, we chose to use *ets* instead of wildtype embryos. *ets* embryos behave similarly to wildtype at early times but do not have a mesoderm invagination (**Extended Data Fig. 1)**, which is a separate complex process of flow and deformation that we do not model explicitly. We first averaged the data over all *ets* embryos where, to decrease embryo-to-embryo variation, we aligned the embryos to the time at which each transitioned to polarized flow (T_asb_ = 0; **Extended Data Fig. 4)**. We performed a fit of the model given by equation (1) to the data with *l*_*H*_, *r*_*a*_, and *r*_*b*_ as free parameters, and found that we could reproduce the symmetric flow (**Fig. 3c**). However, this model could not reproduce the later polarized flow based on the measured myosin profiles (**Fig. 3d, e**) because the spatially averaged velocity is always strictly zero for this simple model (**Fig. 3f**). Non-zero average flow is not possible because equation (1) contains only tissue-intrinsic, symmetric forces and a homogeneous friction force, none of which can lead to polarized flow. Thus, an additional mechanism is required for driving the polarized flow.

### Localized friction or adhesion is not responsible for polarized flow

During the polarized phase of flow, the dorsal-posterior activation of apical myosin (**Fig. 2a, 3d**) leads to bending of the epithelium. This causes the anterior end of the apical myosin domain to come into close contact with the eggshell^29^, which could create a localized domain with higher friction. This close contact could also lead to adhesion between the epithelium and the eggshell due to the presence of the adhesion protein alpha-Integrin (Scab), which is expressed in this region^39^. Intuitively, this configuration could lead to asymmetric contraction of the myosin patch since one end is able to move more freely than the other. Adhesion of the anterior end of the apical myosin domain has even been shown to be crucial to the anterior wave propagation of the endoderm invagination slightly later in development^29^. To test whether the polarized flow that we observe could be created by such an asymmetric friction, we accordingly updated our model by including a domain (*G*) that translocates at the anterior end of the myosin domain, where friction is increased by a factor *g* (see equation (2) in **Fig. 4a)**.

**Figure 4:**
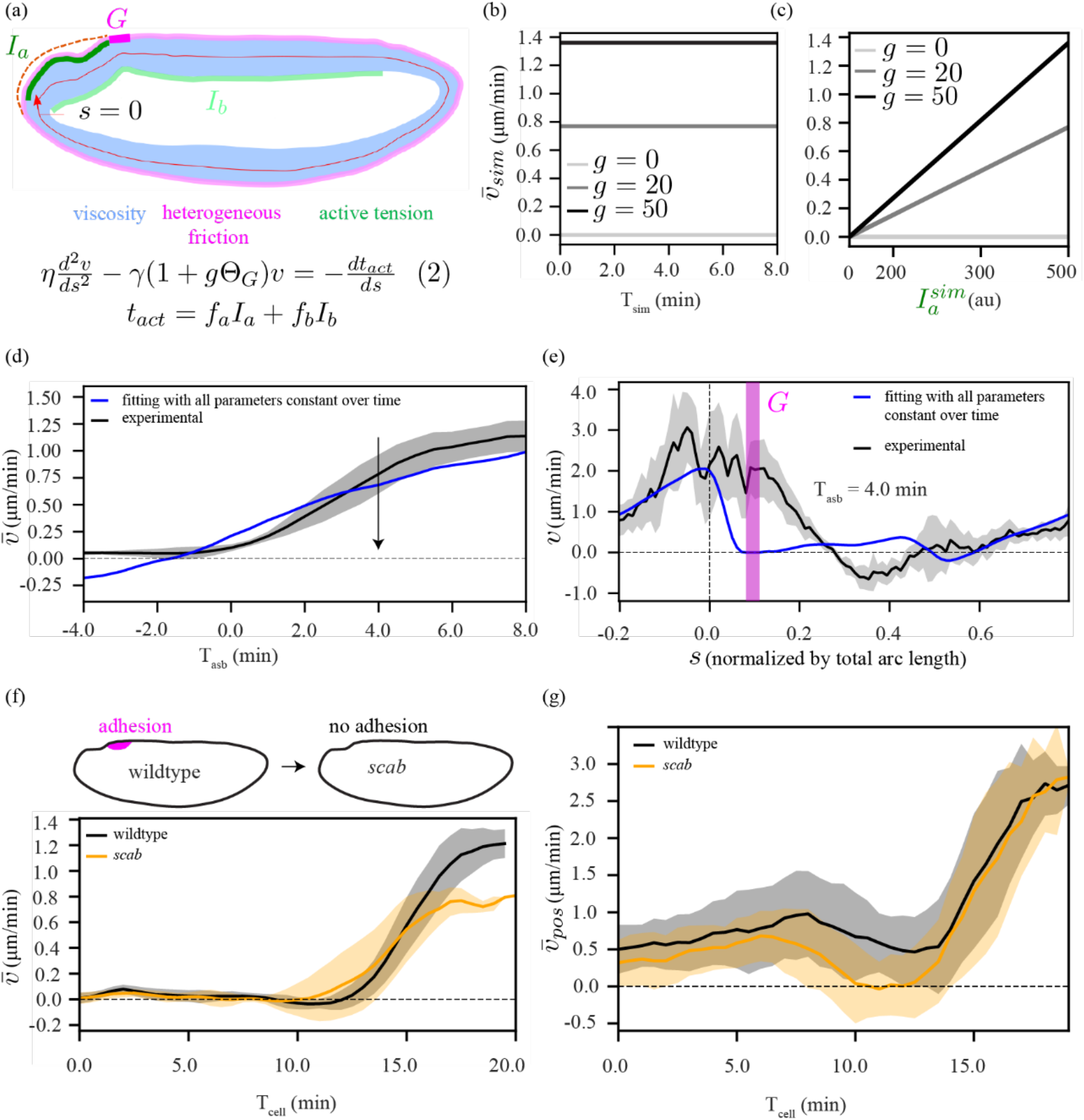
A model coupling adhesion (or heterogeneous friction) with active tension cannot explain the polarized flow. **(a)** Schematic representation of our model, equation (2), which is similar to equation (1) in **Fig. 3a**, but with an additional domain *G* (magenta) where the localized friction is increased by a factor *g*. **(b)** Temporal profile of the spatially averaged velocity 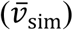 in our simplified simulations (**Extended Data Fig. 5a, b**) when myosin intensity 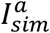 is constant over time, shown for three different values of *g*. **(c)** Dependence of the simulated average velocity 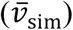 on myosin intensity 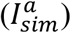 and localized friction increase *g*. **(d)** Experimentally measured temporal profile of spatially averaged velocity 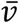 (black) and fit of equation (2) (blue). The fit was performed using the same procedure as described in Fig. 3 (see Supplementary Information). **(e)** Fit curve from the simultaneous fit in panel **d** at a presentative time point during the asymmetric phase (T_asb_ = 4 min). Fitting with a time-dependent localized friction *g* resulted in a similar outcome (**Extended Data Fig. 5c, d**). **(f)** (*top*) schematic diagram of *scab* knockout (*bottom*), temporal profile of spatially averaged velocity 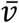 for wildtype (black) and *scab* (orange). **(g)** Temporal profile of spatially averaged velocity 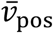, where we averaged only over a posterior domain of the embryo, from *s* = -0.1 to *s* = 0.15. For further details see **Extended Data Fig. 5** and Supplementary Information. In all panels, the shaded regions indicate the standard deviation, computed over 6 embryos.

To build intuition for the effect of this new term in the model, we performed simulations using our elliptic model (**Extended Data Fig. 5a, b**). We obtained polarized flow, though the average velocity remained constant in time unless the amount of localized friction *g* (**Fig. 4b**) or the myosin intensity (**Fig. 4c**) changed over time. The latter could be a possibility as our observed data shows apical myosin increasing over time (**Fig. 2c**).

To test the localized-friction model against the experimental data, we performed detailed fits of the measured velocity to solutions of equation (2) using the measured time-dependent apical and basal myosin patterns, where we included the additional fit parameter *g*, which we restricted to be positive. While we found that this reproduced the observed overall increase in the average velocity (**Fig. 4d**), it failed to predict the observed spatial velocity profile at individual time points (**Fig. 4e**). This is because the model predicts that, for positive values of *g*, the velocity in region *G* should be negative (counterclockwise) or go to zero in the case of infinite friction (as shown using our elliptic model in **Extended Data Fig. 5f, Supplementary Information**), while experimentally we observed positive (clockwise) flow.

We also experimentally tested the contributions of alpha-Integrin-mediated adhesion to polarized flow, by generating a CRISPR knock-out of *scab*^39^ (**Supplementary Video 6**). We found that the movement of the posterior tissue is almost identical for wildtype and *scab* embryos (**Extended Data Fig. 5g**), however the average velocity after symmetry breaking is reduced for *scab* mutants (**Fig. 4f**). To reduce the impacts of any unintended effects of the mutation on other parts of the embryo (**Extended Data Fig. 5h)**, we also computed a spatially averaged velocity over only the posterior domain of the epithelium (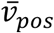; **Fig. 4g**). This analysis confirmed that the onset of polarized flow in *scab* mutant embryos is similar to wildtype, thus indicating that alpha-Integrin is not required for the onset of polarized flow. Taken together, our combined study of modeling and experiment ruled out adhesion (or localized friction) as a mechanism for the onset of polarized flow.

### Interaction of tissue curvature with active moment can explain the polarized flow

Polarized forces, and thus polarized flow, can be created by the interaction of a curvature gradient with an active moment, which follows from the theory of active surfaces^40^ (**Supplementary Information**). Active moments are created when apical and basal myosin tensions differ. When myosin (and therefore active tension) is present at similar levels apically and basally in a region of tissue, this acts to contract the green region, which in turn exerts forces on the surrounding tissue (**Fig. 5a**). When the levels of apical and basal myosin differ in a region of tissue, an active moment is present that causes this region to exert torques on neighboring tissue, which acts to increase or decrease the curvature of the green region (**Fig. 5b**).

**Figure 5:**
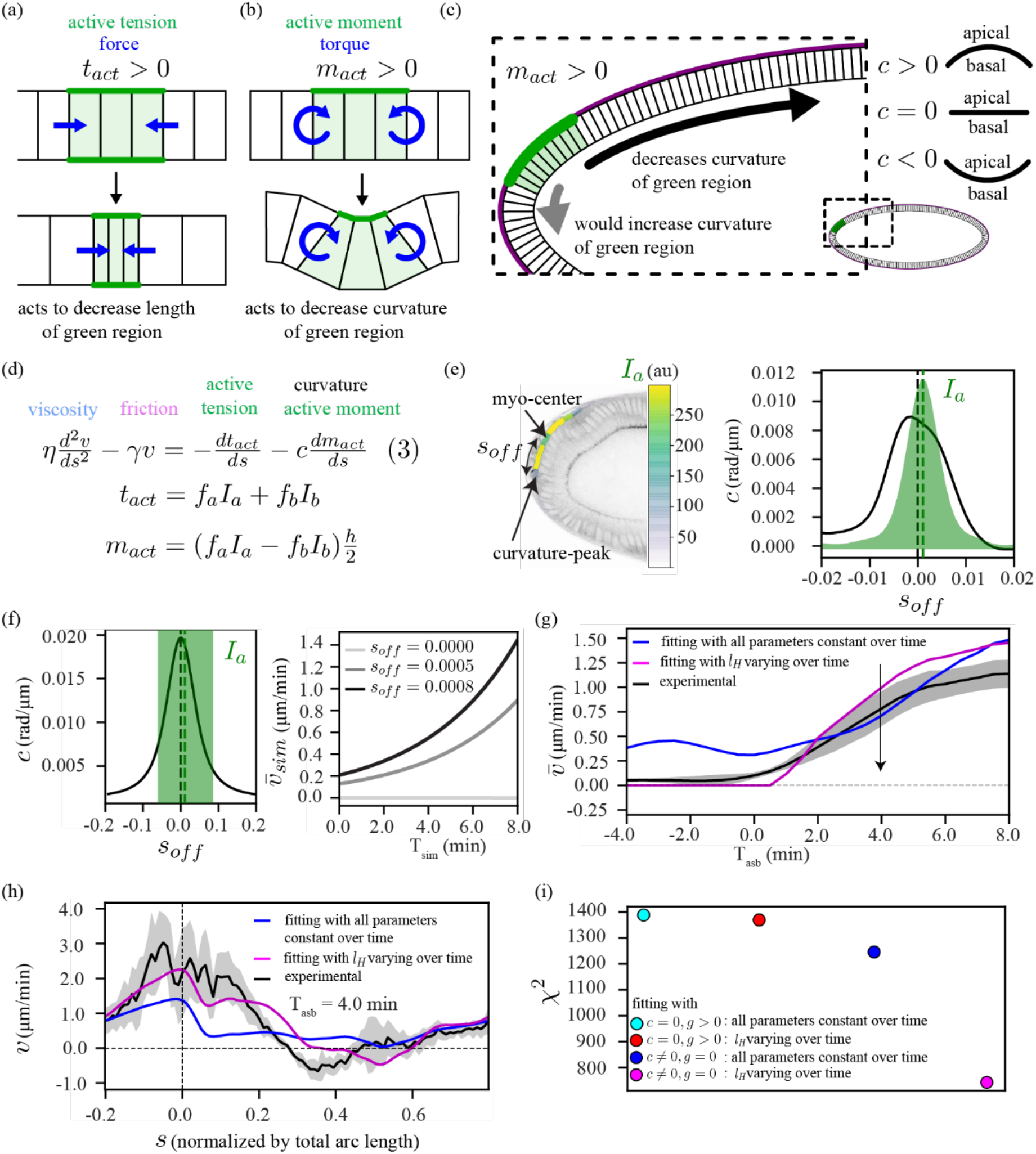
A model including a coupling of tissue curvature with an active moment can explain the polarized flow. **(a-c)** Schematic representation of how a curvature-based mechanism can drive polarized flow: **(a)** active tension (*t*_*act*_) in a region of tissue (green) exerts forces on the surrounding tissue (blue arrows) to contract the green region. **(b)** An active moment (*m*_*act*_) in a region of tissue exerts torques on neighboring tissue (blue arrows) to decrease the curvature of the green region. **(c)** Active moment in some part of the tissue (green) interacts with the shape of the eggshell, where a positive active moment (*m*_*act*_ > 0) drives the tissue in a direction that allows the green region to decrease its curvature (black arrow). Sign convention used: curvature is positive (*c* > 0) where the epithelium is convex, and negative (*c* < 0) where the epithelium is concave. **(d)** Model equation (3): we introduce a new force term to equation (1) that describes a coupling of curvature *c* with active-moment *m*_*a*_. **(e)** Experimental quantification of positional mismatch (*s*_off_) between curvature-peak at the posterior pole and the center of apical myosin domain. **(f**) Simplified simulation using our elliptic model (**Extended Data Fig. 6a, b**). (*left*) illustration of myosin-curvature offset *s*_off_. (*right*) simulated temporal profile of the spatially averaged velocity 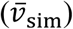 with myosin intensity 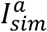 constant over time for three different initial values for *s*_off_. **(g)** Temporal profile of spatially averaged velocity 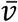 in experiment (black) and two fits (blue and magenta) to equation (3). The fit was performed using the same procedure as described in Fig. 3 (see Supplementary Information). To obtain the fit shown by the blue curve, we set all parameter values, *l*_*H*_, *r*_*a*_, and *r*_*b*_ constant over time (the early polarized flow in this curve is explained in **Extended Data Fig. 6c**). For the fit shown by the magenta curve, we impose constant *r*_*a*_ and *r*_*b*_, while allowing the hydrodynamic length, *l*_*H*_, to be different for each time point (temporal profile of *l*_*H*_ shown in **Extended Data Fig. 6d**). **(h)** Spatial fit curves for velocity, corresponding to the fits in panel **g** for a representative time point during the polarized phase (T_asb_ **=** 4 min). **(i)** Comparison of the fit quality for different hypotheses: chi-square values (*χ*^2^), summed over all time points, for model with locally increased friction, equation (2), with either all parameters constant (cyan) or only varying *l*_*H*_ (red), and model with curvature-active-moment coupling, equation (3), with either all parameters constant (blue) or only varying *l*_*H*_ (magenta). For further details see **Extended Data Fig. 6** and Supplementary Information. In all panels, the shaded regions indicate the standard deviation, computed over 6 embryos.

We therefore hypothesized that polarized flow could arise because the dorsal-posterior apical myosin patch creates an active moment that interacts with the curvature gradient in this region. The emergence of polarized flow can be most clearly explained in the limit where the yolk pressure is so high that it presses the epithelium against the eggshell. In this case, the epithelium is forced to have the same curvature as the eggshell everywhere. Since the region with the apical myosin patch has a positive active moment, it will act to decrease its own curvature, which happens only if it moves away from the pole, creating polarized flow (**Fig. 5c**).

To test whether this effect could explain our observations, we added coupling between the active moment and the epithelium curvature to our quantitative model by including a new force term, as shown in equation (3) (**Fig. 5d, Supplementary Information**). This term follows directly from the theory of active surfaces^40^, and includes no new fit parameters. The resulting flow depends critically on the position of the domain of apical myosin activation, relative to the curvature profile of the embryo. A simple quantity to consider is the offset between the center of the apical myosin domain and the position of peak epithelial curvature at the posterior pole (henceforth called myosin-curvature offset, *s*_off_, **Fig. 5e**). Simulating equation (3) using our elliptic model (**Extended Data Fig. 6a, b**) created polarized flow where the average velocity increased as *s*_off_ increased (**Fig. 5f**) even when we held the myosin intensity constant over time.

To further test our hypothesis, we fit equation (3) to the experimentally measured velocity patterns, using measured myosin and curvature data. Fitting with all physical parameters held constant over time reproduced the substantial increase of the polarized flow with time, but also predicted some polarized flow at times before symmetry breaking (blue curve **Fig. 5g, h**). We found that this early polarized flow in the blue fit curve arises due to the active moment created by basal myosin (**Extended Data Fig. 6c**). This suggested that there must be something that prevents polarized flow at early stages. It has previously been observed that the amount of friction changes over time during the early phase of cellularization in the *Drosophila* embryo^36^. Moreover, tissue deformations and folding that occur during gastrulation alter the distance between epithelium and eggshell in certain regions, which could further affect the friction. Accordingly, we performed fitting allowing the overall homogeneous friction, *γm*, to change over time. This modified fit captured the experimentally observed transition from symmetric to polarized flow and the spatial velocity patterns (magenta curves, **Fig. 5g h**). This fit suggested a large increase of the hydrodynamic length *l*_*H*_ around the time of symmetry breaking, i.e. a large decrease in friction with the eggshell (**Extended Data Fig. 6d**). However, this increase in *l*_*H*_ decreased drastically when we consider that the region expressing apical myosin can only be compressed by a finite amount (**Extended Data Fig. 6e, f, 7, Supplementary Information**). Comparing fitting accuracies using chi-squared analysis, we found that the curvature-active-moment model, equation (3), could consistently explain the measurements better than the tension-with-heterogeneous-friction model, equation (2) (**Fig. 5i**). This supports the idea that coupling between curvature and active moment can act as a driver for the polarized flow observed in the early embryo.

### Changing embryo curvature and active moment alters the flow, consistent with the model

To further test the predictions of the model, we performed targeted perturbations to parameters that are essential for flow in the model: the curvature of the embryo, and the location of the apical myosin domain with respect to the curvature peak (*s*_off_).

To alter the curvature of the embryo, we took advantage of the fact that there is an adhesion molecule, Fat2, present in the maternal follicular epithelium that is needed to shape embryo geometry. We therefore used *UAS-fat2-RNAi, traffic jam(tj)-GAL4* mothers to produce embryos with altered shape^41–43^ (hereafter *fat2* embryos) that are significantly shortened along the anterior-posterior axis and, as a result, are much rounder (**Fig. 6a, b, Supplementary Video 7**). To explore the expected impact of changing the aspect ratio on the flow dynamics, we performed simulations based on equation (3) using our elliptic model with varying aspect ratios. The simulations predicted that decreasing the embryo’s aspect ratio should decrease the rate at which the spatially averaged velocity increases during the phase of polarized flow (**Fig. 6c**). Experimentally, we observed a wide variation in the average velocity profile for individual *fat2* embryos (**Extended Data Fig. 9a**). But, when the embryos were aligned with respect to the onset of polarized flow, we observed the predicted decrease in the average velocity (**Fig. 6d**). This decreased velocity was confirmed by tracking of the position of the posterior tissue (**Fig. 6e**) and by averaging the velocity in only the posterior of the embryo (**Extended Data Fig. 9b**). This result confirms that the geometry of the embryo, plays an important role in the onset and the dynamics of the polarized flow.

**Figure 6:**
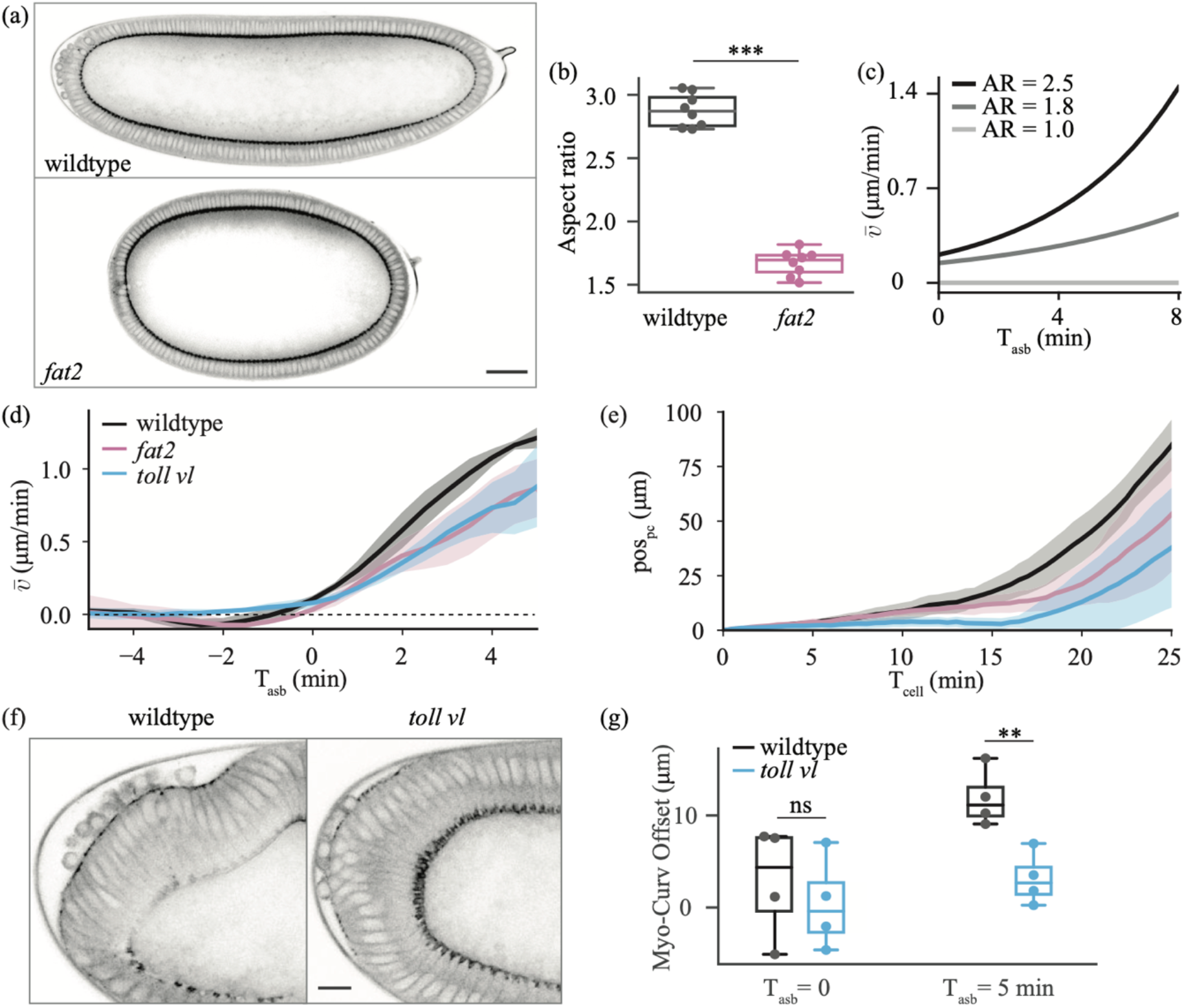
Experimental perturbations to challenge model predictions. **(a)** Sagittal section of wildtype (top) and *fat2* (bottom) embryos at T_cell_ = 0, imaged for *sqh*::GFP. Scale bar is 50 μm. **(b)** Quantification of aspect ratio, defined as the length of the major embryo axis divided by the maximal height of the embryo for 8 wildtype and 8 *fat2* embryos. **(c)** Average velocity of tissue flow resulting from simulations using our elliptic model with different aspect ratios (AR). In the simulations (**Extended Data Fig. 5e**), the position and extent of the myosin domain was initialized consistent with the experimental data. **(d)** Experimental spatially averaged tangential velocity as a function of time after symmetry breaking for 5 wildtype, 4 *fat2*, and 6 *toll vl* embryos. **(e)** Pole cell position (pos_pc_) as a function of time for 6 wildtype, 5 *fat 2*, and 7 *toll vl* embryos. **(f)** View of the posterior of a wildtype (left) and *toll vl* (right) embryo imaged for *sqh*::GFP. Scale bar is 10 μm. **(g)** Quantification of the offset between the position of maximum curvature and the center of the patch of posterior myosin (Myo-Curv Offset) for 4 wildtype and 4 *toll vl* embryos. Comparisons in **b** and **g** performed using two-tailed unpaired t-test. ns, not significant; **, p < 0.01; ***, p < 0.001.

Next, we altered the position of the apical myosin domain using the *toll vl* mutant embryos described previously. Because the *toll vl* mutation removes dorsal-ventral polarity, the domain of apical myosin becomes more centered on the posterior pole of the embryo compared to the dorsal-posterior localization in wildtype (**Fig. 6f, g**). The difference becomes increasingly significant over time. According to our prediction, decreasing the myosin-curvature offset, *s*_off_, is expected to decrease the average velocity (**Fig. 5g**). Once the individual embryos (**Extended Data Fig. 9c**) were aligned with respect to the onset of polarized flow, this decrease in the average velocity was confirmed experimentally (**Fig. 6d, e, Extended Data Fig. 9b**).

In addition to the decrease in average velocity, there is a large delay (approximately 5 min) in the onset of polarized flow in *toll vl* embryos (**Fig. 2e**). This delay likely occurs for two reasons. If the apical myosin patch were perfectly centered on the posterior pole, it would not flow at all until it was displaced slightly in any direction by some other effect. This would also explain why *toll vl* embryos tend to flow in directions other than dorsally as normally seen in wildtype (**Extended Data Fig. 2**). Additionally, *toll vl* embryos lack the dorsal-ventral asymmetry in basal myosin that, in wildtype embryos, leads the myosin patch to shift away from the pole even before the onset of polarized flow. This initial shift increases the speed of polarized flow arising from the myosin-curvature offset in wildtype. There is, in a sense, a feedback based on the initial position of the apical myosin patch and the initial flow. The larger the offset, the more polarized flow, which increases the offset, and so on until the domain of myosin reaches the region of lowest curvature on the dorsal side of the embryo.

We performed two further tests to assess whether the observed change in flow results from the changed position of myosin with respect to the curvature peak. First, we extracted parameters from wildtype fitting (**Extended Data Fig. 10a, b**), and used the myosin and curvature profiles measured in *toll vl* to predict flow using model equation (3). This prediction yielded a good match to the experimentally measured flows in *toll vl* embryos and confirmed the decrease in the average velocity (**Extended Data Fig. 10c, d**). Secondly, we quantified flow dynamics in *capicua* mutant embryos, whose apical myosin domain is larger but more centered on the posterior pole than in wildtype embryos (**Extended Data Fig. 9d, e, Supplementary Video 8**). In *capicua* embryos we observed a slowed and delayed increase in average velocity despite a larger domain of apical myosin (**Extended Data Fig. 9f, g**), which was also predicted by simulations using our elliptic model (**Extended Data Fig. 9h**).

Taken together, these experimental perturbations deeply and broadly challenge the predictions of the model. We therefore conclude that the model given in equation (3) accurately captures the features that are necessary for driving the onset and evolution of polarized flow in early *Drosophila* morphogenesis. Overall, our results show that an initially symmetric flow, driven by a non-uniform pool of basal myosin, transitions to polarized flow as a result of the activation of apical myosin in a domain of the embryo that experiences a curvature gradient.

## Discussion

We have identified two distinct phases of tissue flow in the early *Drosophila* embryo, where an initially symmetrically deforming tissue gradually transits to a polarized flow. While previous work has studied flow-generating mechanisms during morphogenesis, we showed that this polarized flow can neither be explained by forces emerging from other tissues such as the germband and mesoderm (**Extended Data Fig. 1**), nor by a myosin-adhesion coupling. Our work revealed instead that a coupling between tissue curvature and an active moment, generated by a difference in apical and basal myosin intensity, is responsible for driving the polarized flow. We also demonstrated the importance of spatially varying curvature, with an offset between the curvature peak and the domain of the active moment for early polarized movement.

This system is an ideal example for how organized multicellular dynamics that occur during morphogenesis emerge from inherited genetic, and geometric blueprints inherited from the mother. Beyond simply fulfilling inherited instructions, the offset between the patterns of cell contractility and tissue curvature is also amplified by the very flow it induces. This illustrates that interactions between genetic and geometric information update this information as a result of morphogenesis. Thus, information is not simply inherited, it is constantly modified.

While the mechanism that we report accurately describes the onset of polarized flow, it is not sufficient to explain later stages of flow, as the decreased curvature gradient towards the dorsal side of the embryo is unfavorable for sustained flow. Indeed, it has been shown that later phases of flow do depend on the process of germband elongation, driven by myosin polarization in the lateral ectoderm^14–16^, and on dynamically regulated adhesion between the epithelium and the vitelline membrane in the dorsal posterior^29^.

To keep our model as simple as possible while still revealing the mechanisms required to achieve the observed flow, we restricted our investigation to the 1D tangential flow in the sagittal plane of the epithelium and made several simplifying assumptions. Quantitative comparisons of the model to our data suggested that contraction of the tissue in the primordium is limited by elastic resistance, which for simplicity, we modeled by a local increase in viscosity (**Extended Data Fig. 7**). This is consistent with the fact that, due to the incompressibility of the cytoplasm, there is a limit to cell deformation upon contraction (**Supplementary Video 1)**. However, for a complete understanding of tissue flow, deformation, and folding, explicit consideration of elasticity and a full 3D modeling approach will be necessary. This would open the way to addressing for example how tissue flow progresses along the dorsal midline during later stages of embryo development.

Morphogenetic processes are both self-organized and dependent upon initial conditions to deterministically guide future processes^2^. In *in vitro* synthetic systems, such initial conditions are engineered to drive robust organoid development^44,45^, but *in vivo* they are inherited from previous developmental stages. During development, heredity is classically associated with genomic heredity. Yet, as we show here, structural or geometric heredity (eg. egg size and shape) is also essential to drive tissue flow and morphogenesis. It will be interesting to investigate further how the interplay between genetic and geometric heredity guide developmental processes in other systems.

## Methods

### Fly strains and genetics

The following mutant alleles were used: *eve*^3^ (Bloomington stock 299), *twi*^1^ (Bloomington stock 2381), *sna*^18^ (Bloomington stock 2311), *toll*^rm9^ and *toll*^rm10^ (gift from Maria Leptin), *cta*^RC10^ (gift from Maria Leptin), *cic*^1^ (gift from Gerardo Jiménez), *traffic jam (tj)-GAL4* (*P{w[+mW*.*hs]=GawB}NP1624 / CyO, P{w[-]=UAS-lacZ*.*UW14}UW14*) (Kyoto Stock Center 104055) and *UAS-fat2 RNAi* (*P{GD14442}v27113*) (Vienna Drosophila Resource Center 27113) (gifts from Sally Horne-Badovinac), scabKO (generated in the laboratory using CRISPR by Jean-Marc Philippe), *osk-Gal4, UASp-CIBN-pmGFP*, and *UASp-CRY2-RhoGEF2* (gift from Stefano de Renzis). The triple mutant ;*eve*^3^, *twist*^1^, *snail*^18^; used was generated in the laboratory by Claudio Collinet.

Myosin regulatory light chain (MRLC) is encoded by the gene *spaghetti squash* (*sqh*, Genebank ID: AY122159). Live imaging of *sqh* was performed using *sqh-sqh::GFP* (on chromosome 2, and on chromosome 3, gift from Robert Karess). Live imaging of the plasma membrane was carried out using *sqh-GAP43::mScarlet* (on chromosome 2 (9736, 2R, 53B2) and 3 (9744, 3R, 89E11) made in the laboratory by Jean-Marc Philippe). The recombinants ;*sqh-sqh::GFP,sqh-GAP43::mScarlet;* and *;;sqh-sqh::GFP,sqh-GAP43::mScarlet* were generated in the laboratory. All unique fly lines generated for this study are available from the corresponding authors upon reasonable request.

Crosses for *toll vl*: virgin *;sqh-sqh::GFP,sqh-GAP43::mSc;toll*^*rm9*^*/TM6C* females were crossed with *;sqh-sqh::GFP,sqh-GAP43::mSc;toll*^rm10^*/TM6C* males. Homozygous offspring were put in a cage.

Crosses for *fat2*: virgin ;*tj-Gal4;sqh-sqh::GFP,sqh-GAP43::mSc* females were crossed with ;*UAS-fat2 RNAi;sqh-sqh::GFP,sqh-GAP43::mSc* males. F1 virgins were crossed with ;*UAS-fat2 RNAi;sqh-sqh::GFP,sqh-GAP43::mSc* males. Resulting progeny were put in a cage.

### Sample preparation

Flies were kept in a cage at 25°C with the exception of GAL4 lines which were kept at 18°C. Embryos were collected using apple cider plates smeared with yeast paste. Embryos were transferred to a mesh metal basket, rinsed with water, dechorionated with 2.6% bleach for 1 min, then rinsed copiously with water before being transferred back to clean agar. Embryos in the early stages of cellularization were visually selected, and aligned laterally. They were then transferred to a glass coverslip coated with a homemade glue. A drop of Halocarbon 200 Oil (Polysciences; for DIC experiments) or 1x phosphate buffered saline (prepared from Dulbecco’s phosphate buffered saline (eurobio); for two photon experiments) was placed on the embryos to keep them from drying during imaging.

### Brightfield imaging

For quantifying the direction of tissue rotation in *toll vl* mutants (**Extended Data Fig. 2c**), embryos were imaged at 21°C on a Zeiss Axiovert 200M inverted microscope using a 20x-0.75 numerical aperture (NA) objective. Images were acquired once per minute for 2-3 hours.

### Two photon imaging

For all other experiments, embryos were imaged at 23.5-24.5°C using a Nikon A1R MP+ multiphoton microscope with a 25x-1.10 NA water-immersion objective. Illumination was provided by a pulsed 1040 nm laser (Coherent) for mScarlet and tunable wavelength pulsed laser (Coherent) set to 920 nm for GFP. Images were acquired sequentially once every 30 seconds with 2x line averaging.

### Data Analysis

#### Pole cell tracking

We selected a pole cell in the center of the cluster and manually tracked its coordinates over time in FIJI. As the selected pole cell began to invaginate, we tracked the position radially outwards from it near the vitelline membrane to record only its tangential movement (**Supplementary Video 3**). The position is defined as 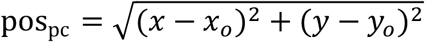 where (*x*_*o*_,*y*_*o*_) is the position at T_cell_ = 0.

#### Image analysis

For both membrane and myosin channels, we used ImageJ software to access images of the sagittal section of the embryo from relevant time points by splitting a corresponding time series movie into individual time frames. For extracting the apical and basal contour of the epithelium, we manually segmented a large number of frames to precisely define the respective apical and basal contours. The separation of the apical contour from the vitelline membrane is a non-trivial segmentation task, we therefore used these initially segmented frames to train a Deep Learning algorithm (namely a U-NET^46^) to do similar segmentation automatically for future movies.

To align the embryos in space, an ellipse (**Fig. 1c** dashed blue ellipse) with direction of the principal axis (dashed blue line) towards the posterior side, was fitted on the apical contour of the epithelium at T_cell_ = 0 min (reference time when cellular front of epithelial cells passes the nucleus). The intersection (indicated by green star) of the principal axis with the midline (in red) of the epithelium is defined as the “zero (*s* = 0)” reference of the arc-length coordinate.

For myosin quantification, we developed a python script to identify the segmented apical and basal contours as closed polynomials. These polynomials were then discretized by 100 evenly spaced nodes, such that each apical node has a correspondence to the nearest basal node. For each node, a myosin-mask was defined by a quadrilateral with height approximately 10 pixels (determined by the thickness of the myosin signal) and variable width determined by the distance between the adjacent nodes. Myosin intensity at a given node was calculated by averaging the pixel intensities within the respective mask.

To extract model inputs, we constructed a midline contour by a new set of nodes defined by the average of each pair of respective apical and basal nodes. At each midline node, we computed tissue velocity via particle image velocimetry (PIV, using python library openpiv), total myosin intensity (sum of the myosin intensity at the apical and basal nodes), active-moment (product of the difference in myosin intensity at the apical and basal nodes with the distance of the midline node from either the apical or the basal node) and curvature (spatial derivative of the angle between the adjacent pair of midline edges).

### Modeling and simulations

Full details on the modeling and the simulations can be found in the Supplementary Information.

## Data availability

The datasets generated during and/or analyzed during the current study are available from the corresponding author on reasonable request.

## Code availability

The computer code used to perform data analysis, simulations, and model fitting is available from the corresponding author on reasonable request.

## Acknowledgements

We thank all members of the Lecuit group for useful discussions. We thank Marc-Eric Perrin for help with machine learning used for embryo segmentation; Claudio Collinet for help in conceiving the project and for preliminary experiments and fly work; and Jean-Marc Philippe, Elise da Silva, and Maxime Louis for designing and generating the molecular constructs used. We thank the Bloomington for providing fly stocks and the IBDM animal facility. The IBDM imaging platform and the France-BioImaging infrastructure supported by the Agence Nationale de la Recherche (ANR-10-INSB-04-01 ; call “Investissements d’Avenir”) provided support. E.W.G. and B.C. were supported by the ERC Advanced Grant SelfControl 788308 awarded to T.L. T.L. is supported by the Collège de France. M.M. thanks the Centre Interdisciplinaire de Nanoscience de Marseille (CINaM) for providing office space. M.M. received funding from the Turing Center for Living Systems (CENTURI), funded by France 2030, the French Government program managed by the French National Research Agency (ANR-16-CONV-0001), and from Excellence Initiative of Aix-Marseille University - A*MIDEX.

## Author contributions

E.W.G., B.C., M.M. and T.L. conceived the project. E.W.G. and T.L. planned the experiments. E.W.G. performed the experiments. B.C. developed the model, performed simulations, and data fitting with the help of M.M. E.W.G. and B.C. performed all data analysis. E.W.G., B.C., M.M. and T.L. discussed the data and wrote the manuscript.

## Competing interest declaration

The authors declare no competing interests.

## Additional information

Supplementary Information is available for this paper.

Correspondence and requests for materials should be addressed to T.L. or M.M.

## Extended Data Figures

**Extended Data Figure 1:**
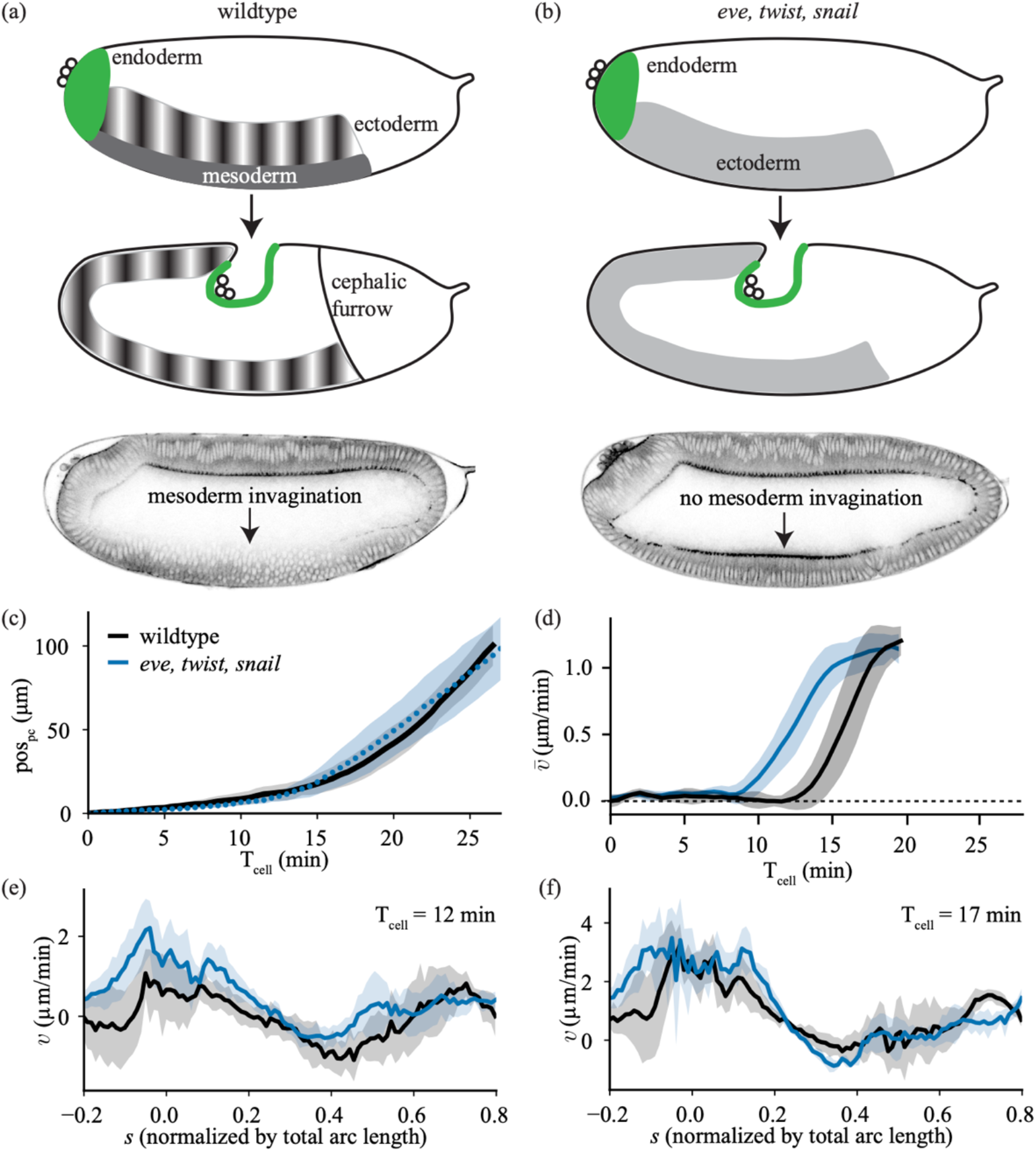
Comparison of wildtype with *eve, twist, snail* mutant embryos. **(a, b)** Cartoons of *Drosophila* embryo (*top*) at an early stage, during the process of cellularization and (*middle*) approximately 30 minutes later for a (**a**) wildtype and (**b**) *eve, twist, snail* (*ets*) mutant embryo. This shows that in *ets* mutants the mesoderm is no longer specified, there is no planar polarization of myosin in the ectoderm, and there is no formation of the cephalic furrow. (*bottom*) Images of these embryos at T_cell_ = 19 min. (**c**) Quantification of the position of the pole cells (pos_pc_) as a function of time since the cellularization front passes the nuclei in the dorsal posterior (T_cell_). Average performed over 6 wildtype and 7 *ets* embryos. (**d**) Spatial average of the tangential velocity as a function of time. Average performed over 5 wildtype and 6 *ets* embryos. (**e, f**) Spatial profile of tangential velocity for wildtype and *ets* embryos at (**e**) T_cell_ = 12 min and (**f**) T_cell_ = 17 min. Error bars represent standard deviation.

**Extended Data Figure 2:**
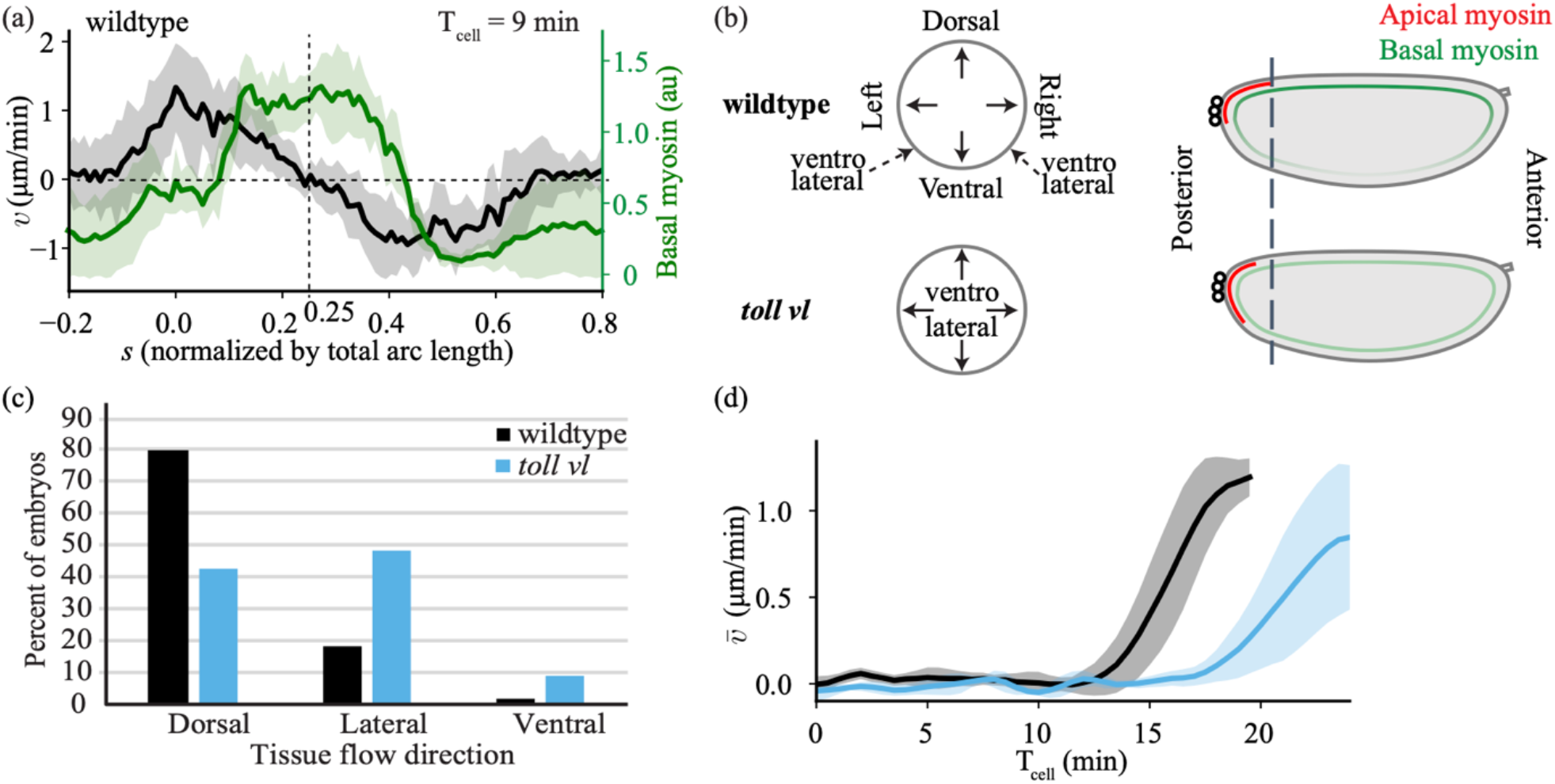
Characterization of *toll vl* embryos. (**a**) Spatial profile of tangential velocity and basal myosin intensity for wildtype embryos at T_cell_ = 9 min. Vertical dashed line represents the center of the dorsal side of the embryo (*s* = 0.25). (**b**) Schema of the difference between wildtype and *toll vl* mutant embryos shown in (*left*) a cross section along the anterior-posterior axis and (*right*) in a sagittal plane. (**c**) Quantification of the direction of tissue flow in wildtype and *toll vl* mutant embryos. Dorsal and ventral indicate that the tissue flows in the imaging plane either towards the dorsal or ventral side and lateral refers to any embryo where the tissue flow occurred out of plan. See **Supplementary Video 4** for examples of each. Data was collected on a DIC microscope for 58 wildtype embryos and 68 *toll vl* mutant embryos (see Methods). A Fisher’s exact test was used to compare dorsal vs non-dorsal flow outcomes between wildtype and *toll vl* conditions yielding a p-value < 0.0001. (**d**) Spatial average of the tangential velocity as a function of time for wildtype and *toll vl* embryos as in **Fig. 2e**, but including later times. Error bars represent the standard deviation.

**Extended Data Figure 3:**
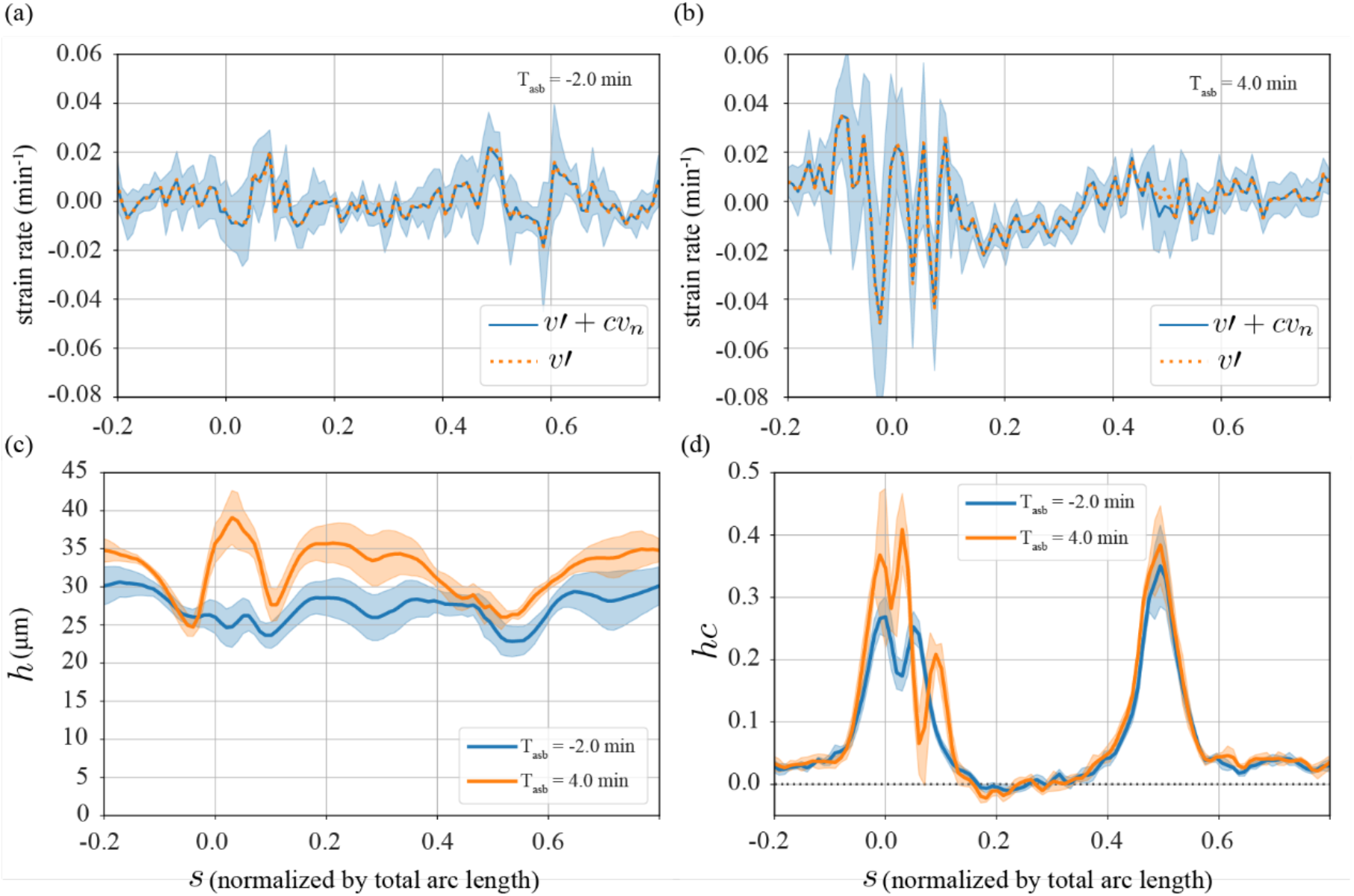
Orders of magnitude in the experimental data. Data from *eve, twist, snail* mutants, which behave similar to wildtype at early times, but which does not show mesoderm invagination, so the height *h* can be measured more accurately. **(a**,**b)** The local strain rate (blue solid curve) is virtually identical with the derivative of the tangential velocity *v′* (orange dashed curve) both at **(a)** T_asb_ = -2 min and **(b)** T_asb_ = 4 min. Thus, the contribution by the normal motion of the epithelium, *v*_*n*_ is negligible. **(c)** The fluctuations in epithelial height *h*(*s*) are on the order of 10% (spatial coefficient of variation). **(d)** The product *hc* is larger at the poles of the embryo, where it maximally becomes approximately 0.4. In all panels, the shaded regions indicate the standard error of the mean, computed over 6 embryos.

**Extended Data Figure 4:**
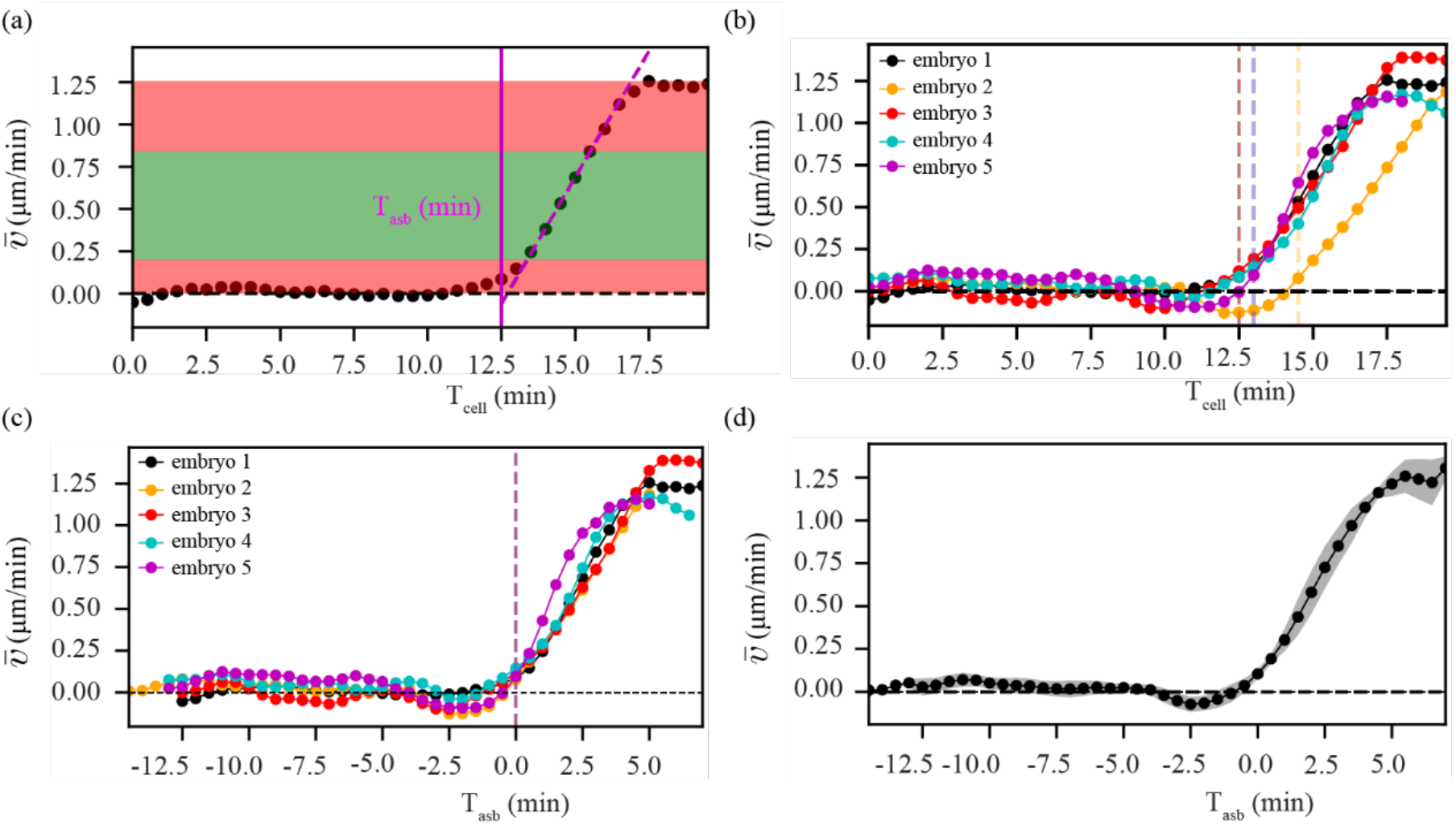
Detection of symmetric to polarized transition in flow. **(a)** Temporal profile of the spatially averaged velocity 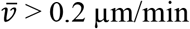 (black dotted curve) computed from the velocity field of individual time frames. A line is fitted in the green region (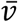 > 0.2 μm/min and the next five time points) to get an intercept with the time axis. This time of intercept becomes the time of symmetry breaking T_asb_ = 0 (vertical magenta line) used to align different embryos. **(b)** Temporal profile of 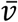 with detection of T_asb_ for many embryos, using the method described in **a. (c)** Temporal profile of 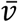 with rescaled time axis, where T_asb_ of the respective embryos is defined as zero reference, i.e T_asb_ < 0 min correspond to symmetric phase of flow and T_asb_ > 0 corresponds to polarized phase of flow. **(d)** Temporal profile of 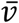, now averaged over all the embryos shown in **c**. The shaded region indicates the standard deviation, computed over 5 embryos.

**Extended Data Figure 5:**
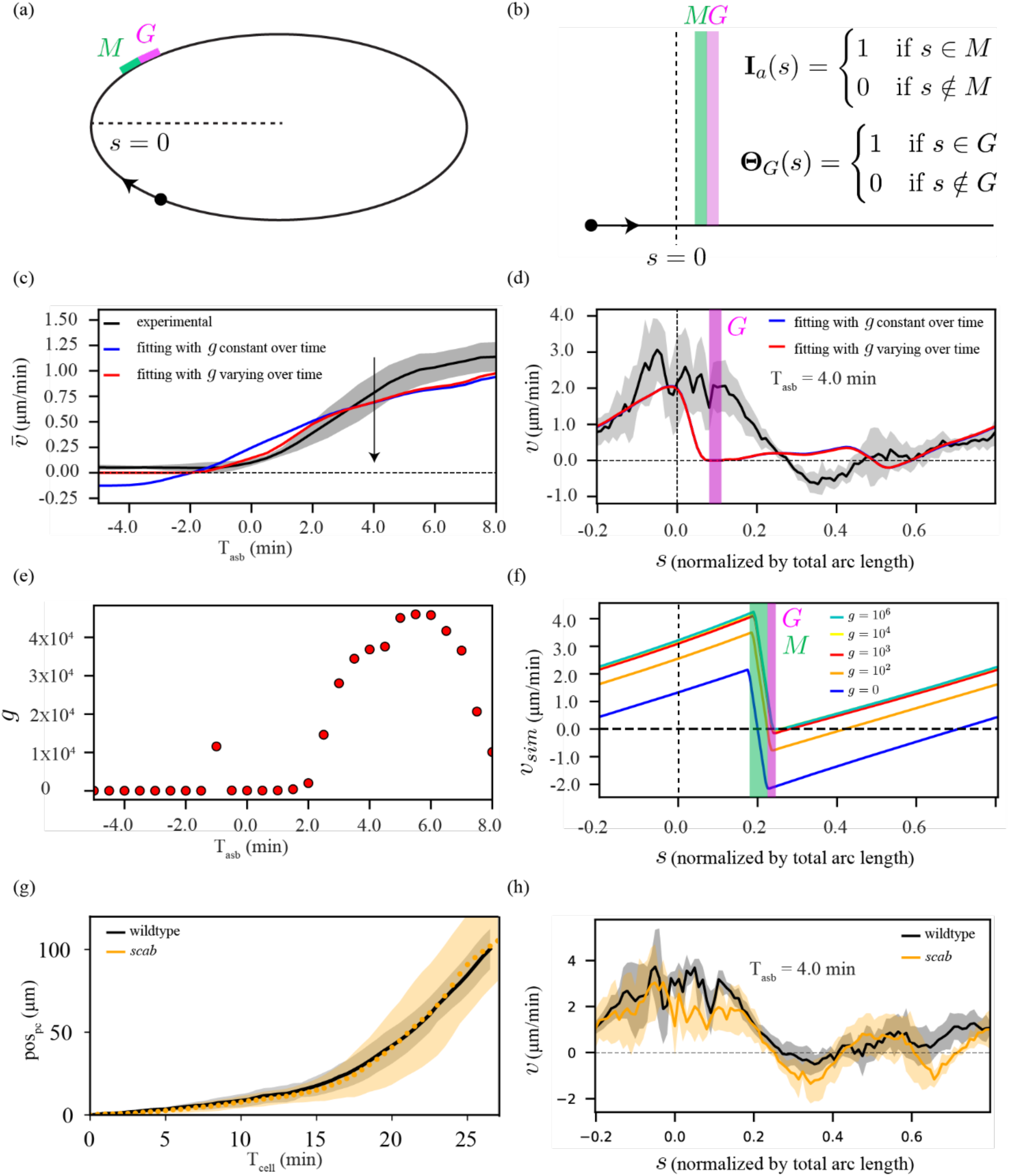
Fitting with heterogeneous friction. **(a)** Schematic of the elliptic representation of the embryo. Green region corresponds to domain of myosin and magenta region corresponds to domain of high friction. **(b)** 1D flat representation of **a**, where the domain of myosin M and domain of high friction G, both are mathematically described by rectangular functions. **(c)** Experimentally measured temporal profile of spatially averaged velocity 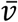 (black) and result of a fit of equation (2). Two fits were performed using the same procedure as described in **Fig. 3** with: (1) all parameter values, *g, l*_*H*_, *r*_*a*_, and *r*_*b*_ constant over time (blue) and (2) constant *r*_*a*_ and *r*_*b*_, while allowing *g* to be different for each time point (red). **(d)** Spatial fit curves for velocity, corresponding to the fits in panel **c** for a representative time point during the polarized phase (T_asb_ **=** 4 min). **(e)** Temporal profile of *g* corresponding to the red curves in panel **c** and **d. (f)** Retrograde flow in the region of high friction *G*, when simulating equation (2) using elliptic embryo in panel **a**. Simulated spatial profile of velocity (*v*_sim_) for varying values *g*. (**g**) Quantification of the position of the pole cells (pos_pc_, see Methods) as a function of T_cell_. Average performed over 6 wildtype (black) and 6 *scab* (orange) embryos. (**h**) Experimentally measured spatial velocity profile (*v*) in wildtype (black) and *scab* embryos (orange) at a representative time point T_asb_ = 4 min. Average performed over 5 wildtype and 5 *scab* embryos. In all panels, the shaded regions indicate the standard deviation.

**Extended Data Figure 6:**
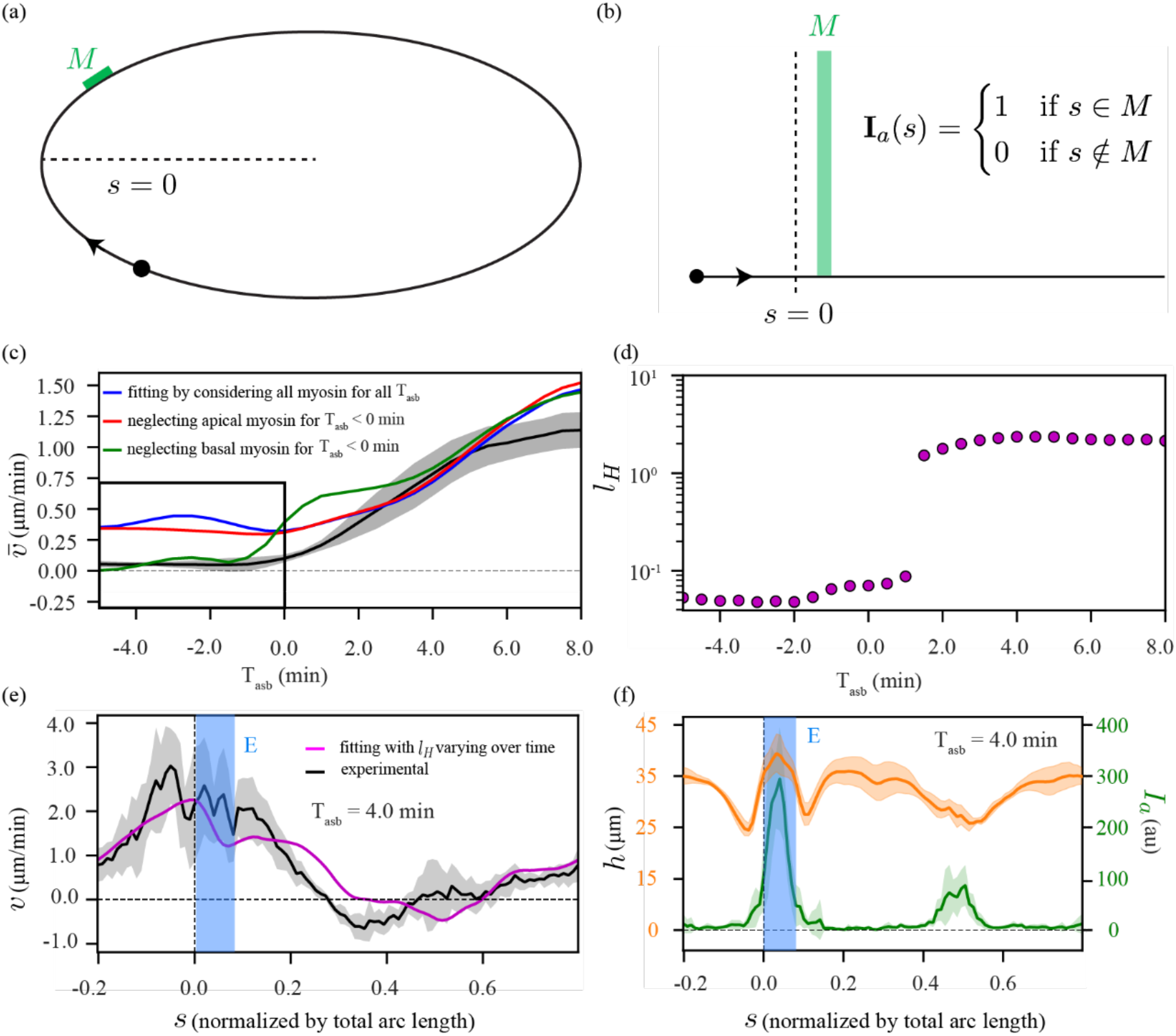
Fitting with curvature-moment coupling. **(a)** Schematic of the elliptic representation of the embryo. Green region corresponds to myosin domain. **(b)** 1D flat representation of **a**, where the domain of myosin *M* is mathematically described by a rectangular function. **(c)** Experimentally measured temporal profile of spatially averaged velocity 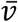 (black) and result of a fit of equation (3). Three fits were performed using the same procedure as described in Fig. 3: (1) considering both apical and basal myosin for the entire time range (blue), (2) neglecting apical myosin for T_asb_ < 0 min (red) and (3) neglecting basal myosin for T_asb_ < 0 min (green). In fitting, we set all parameter values: *l*_*H*_, *r*_*a*_, and *r*_*b*_ constant over time. **(d)** Temporal profile of the hydrodynamic length (*l*_*H*_) corresponding to the magenta curves in **Fig. 5g, h**. In this plot *l*_*H*_ is given in units of epithelial length L=1000 μm. **(e, f)** Understanding the large increase of *l*_*H*_ around the time of symmetry breaking (T_asb_ = 0 min). **(e)** Experimentally measured spatial profile of velocity *v* (black) and the associated fit curve shown in **Fig. 5h** (magenta). **(f)** Experimentally measured spatial profile of the epithelial height (*h*, orange) and apical myosin intensity (*I*_*a*_, green) at same time point as in **e**. In all panels, the shaded regions indicate the standard deviation, computed over 6 embryos.

**Extended Data Figure 7:**
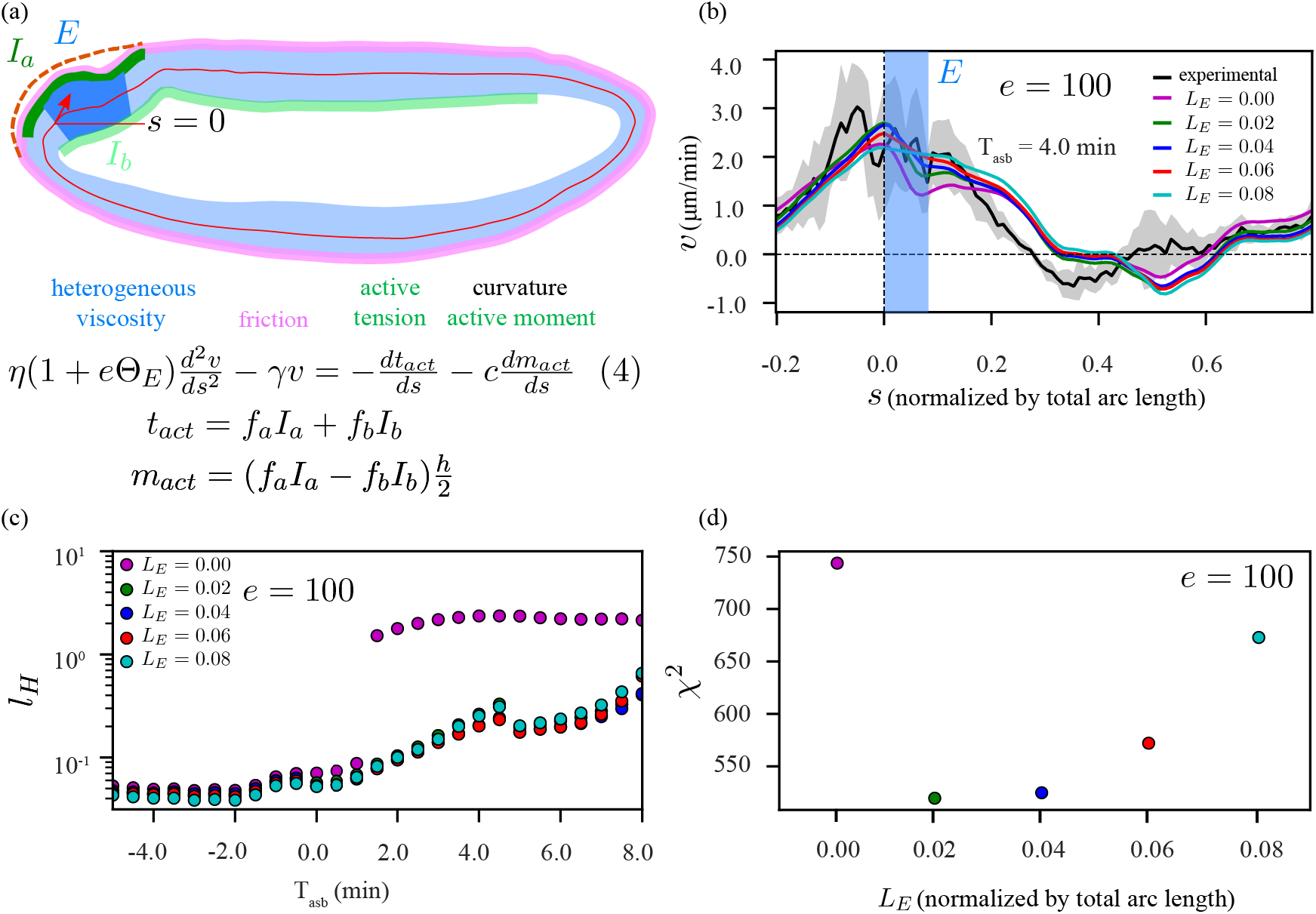
Model with limited tissue contraction at the region of apical myosin. **(a)** Schematic representation of our model, equation (4), which is similar to **Fig. 5d**, but with an additional domain *E* (dark blue region) where the localized viscosity is increased by a factor *e*. **(b)** Experimentally measured spatial profile of velocity 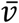(black) and result of fits of equation (4) in panel **a**. Five fit curves, each for a unique choice of the length of high viscosity domain *L*_*E*_ (centered around and restricted within the apical myosin domain, and with increased value of viscosity fixed at *e* = 100). The fits were performed using the same procedure as described in Fig. 3 except that we imposed constant *r*_*a*_ and *r*_*b*_, while allowing the hydrodynamic length, *l*_*H*_, to be different for each time point. **(c)** Temporal profile of *l*_*H*_ corresponding to fitting described in **b. (d)** Comparison of the fit quality for different fitting curves shown in **b**: chi-square values (*χ*^2^), summed over all time points. Here, *l*_*H*_ is given in units of epithelial length *L*∼1000 μm. In all panels, the shaded regions indicate the standard deviation, computed over 6 embryos.

**Extended Data Figure 8:**
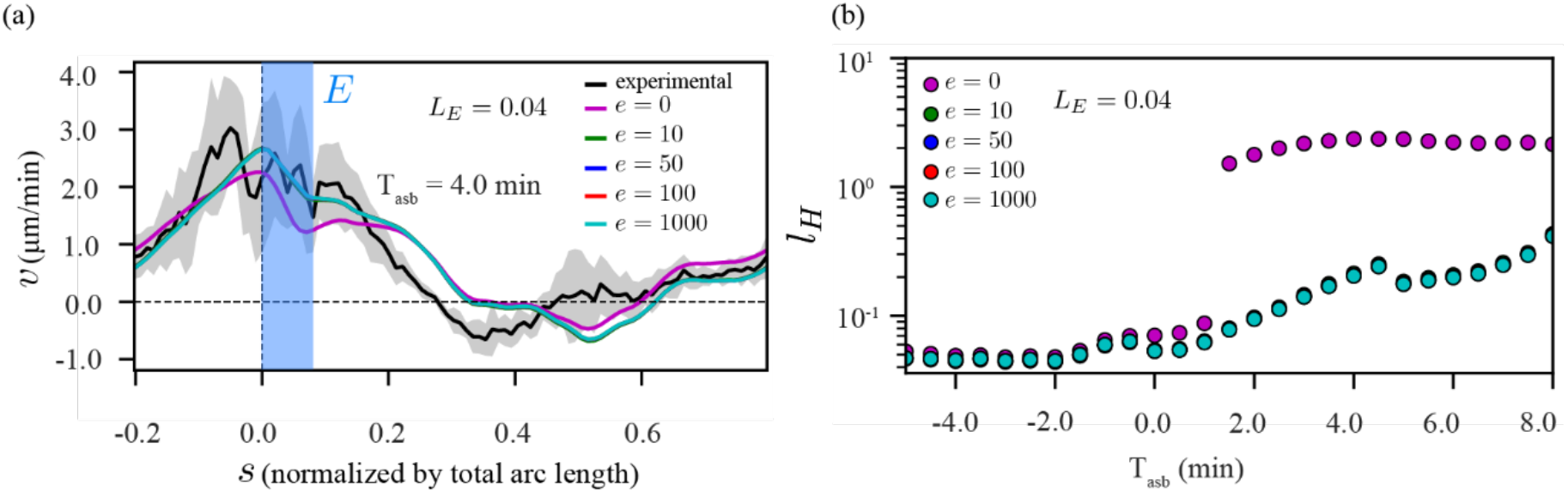
Robustness of fitting with respect to the degree of limited tissue contraction at the region of apical myosin. **(a)** Experimentally measured spatial profile of velocity 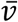 (black) and result of fits of equation (4) in panel **Extended Data Figure 7a**. Five fit curves, each for a unique choice of the values of increased viscosity *e*, where the patch of high viscosity domain (blue region, *E*) was chosen to be of fixed length *L*_*E*_ = 0.04 which falls within and centered around the apical myosin domain. The fit was performed using the same procedure as described in **Extended Data Fig. 7. (c)** Temporal profile of *l*_*H*_ corresponding to fitting described in **a**. Here, *l*_*H*_ is given in units of epithelial length *L*∼1000 μm. In all panels, the shaded regions indicate the standard deviation, computed over 6 embryos.

**Extended Data Figure 9:**
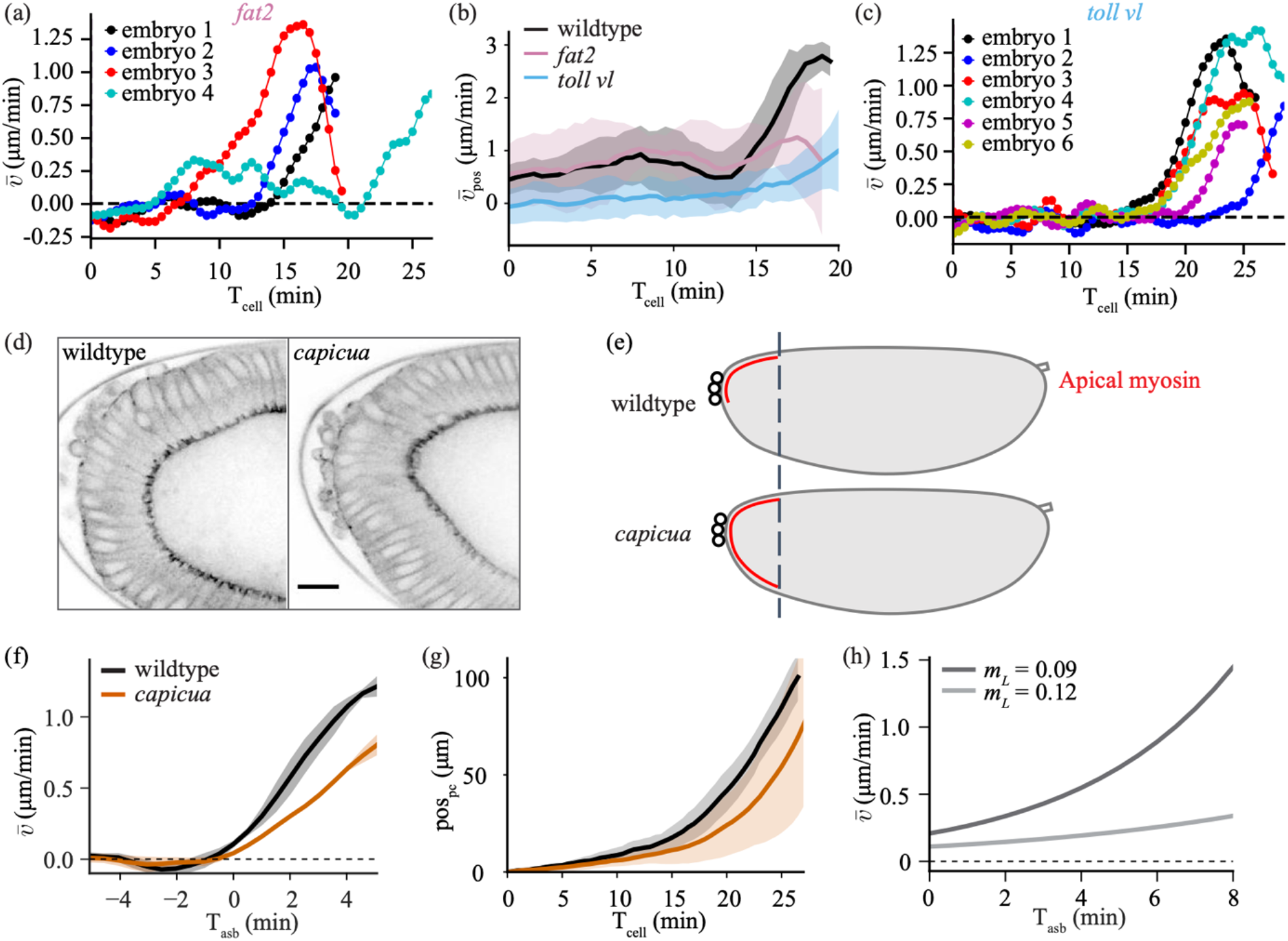
*fat2* and *capicua* characterization. **(a)** Temporal profile of 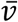 for individual *fat2* embryos. **(b)** Temporal profile of posterior averaged velocity *v*_pos_ for wildtype (black), *fat2* (pink), and *toll vl* (blue) embryos. **(c)** Temporal profile of 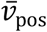 for individual *toll vl* embryos. **(d)** View of the posterior of a wildtype (left) and *capicua* (right) embryo imaged for *sqh*::GFP. Scale bar is 20 μm. Note: the *capicua* embryos were imaged with a single copy of s*qh*::GFP and GAP43::mSc. **(e)** Schematic showing the change in apical myosin domain in *capicua* embryos. **(f)** Experimental spatially averaged tangential velocity as a function of time since symmetry breaking for 5 wildtype, and 5 *capicua* embryos. **(g)** Pole cell position (pos_pc_) as a function of time for 6 wildtype, and 9 *capicua* embryos. **(h)** Average velocity of tissue flow resulting from simulations performed on elliptical embryos with different length of myosin domain (*m*_*L*_; see **Supplementary Information**).

**Extended Data Figure 10:**
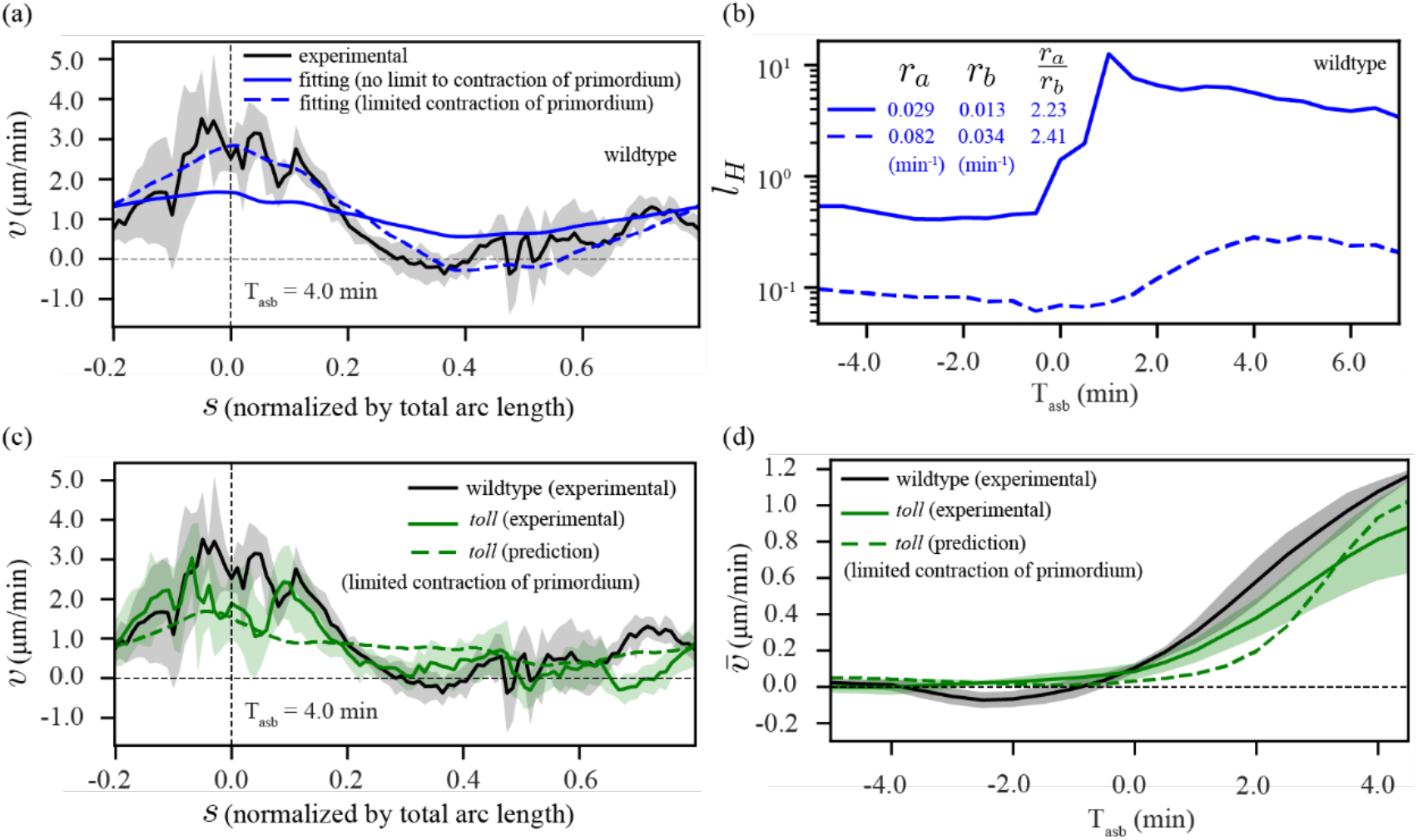
wildtype, *toll vl*, fitting and prediction. **(a)** Experimentally measured spatial profile of velocity *v* (black) in wildtype embryos and result of fits of equation (3) in panel **Fig. 5d** (blue solid curve, no limit to contraction of the primordium) and equation (4) in panel **Extended Data Fig. 7a** (blue dashed curve, limited contraction of the primordium with *e* = 100 and *L*_*E*_ = 0.04), representative time T_asb_ = 4 min. The fit was performed using the same procedure as described in **Fig. 5g** (magenta curve). **(b)** Parameters corresponding to fits in panel **a**, temporal profile of the hydrodynamic length *l*_*H*_ (as curves) and other constant parameters *r*_*a*_, *r*_*b*_ and their ratio (in legend). **(c, d)** Prediction of velocity for *toll vl* using parameters from wildtype in panel **b** (blue dashed line), and myosin and curvature data from *toll vl*. **(c)** Experimentally measured spatial profile of velocity v in wildtype (black) and *toll vl* (green solid curve), and predicted spatial velocity profile for *toll vl* (green dashed curve). **(d)** Experimentally measured temporal profile of spatially averaged velocity 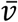 in wildtype (black) and *toll vl* (green solid curve), and predicted temporal profile of spatially averaged velocity for *toll vl* (green dashed curve).

## Supplementary Videos

**Video 1**: Time-lapse of early *Drosophila* morphogenesis in a wildtype embryo. Imaged in the sagittal plane on a two-photon microscope labeled with GAP43:mScarlet (top) and *sqh*:GFP (bottom).

**Video 2**: Tissue dynamics in wildtype and *ets* embryos. Time-lapse of myosin activation in a wildtype (top) and *eve, twist, snail embryo* (bottom) synchronized with respect to the time when the cellularization front passes the nuclei in the dorsal posterior.

**Video 3**: Tracking of pole cells in a wildtype embryo. Time-lapse of tissue dynamics in an embryo labeled for cell membrane maker GAP43:mScarlet. The green dot shows the position used to calculate pole cell movement over time (as in **Fig. 1d**).

**Video 4**: Direction of flow in *toll vl* embryos. Time-lapse of three *toll vl* mutant embryos that flow indifferent directions. The top embryo flows dorsally, the middle embryo flows laterally, and the bottom embryo flows ventrally.

**Video 5**: Tissue dynamics in wildtype, *toll vl*, and *cta* embryos. Time-lapse of myosin activation in a wildtype (top), *toll vl* (middle), and *cta* (bottom) embryos synchronized with respect to the time when the cellularization front passes the nuclei in the dorsal posterior.

**Video 6**: Tissue dynamics in wildtype and *scab* knockout embryos. Time-lapse of myosin activation in a wildtype (top) and *scab* knockout (bottom) embryos synchronized with respect to the time when the cellularization front passes the nuclei in the dorsal posterior.

**Video 7**: Tissue dynamics in wildtype and *fat2* embryos. Time-lapse of myosin activation in a wildtype (top) and *fat2* (bottom) embryos synchronized with respect to the time when the cellularization front passes the nuclei in the dorsal posterior.

**Video 8**: Tissue dynamics in wildtype and *cic* embryos. Time-lapse of myosin activation in a wildtype (top), and *cic* (bottom) embryos synchronized with respect to the time when the cellularization front passes the nuclei in the dorsal posterior.

## Notes

### Competing Interest Statement

The authors have declared no competing interest.

