## Supplementary Information for "Genetic and geometric heredity interact to drive polarized flow in the *Drosophila* embryo"

A Force and torque balance 12

B Virtual work by external forces and torques 13

### 1 Derivation of the model equation

In this section we derive the model equation Eq. (3) in the main text that predicts the tangential flow in the embryo. In brief, this equation describes the embryo as an overdamped liquid (Extended Data Fig. 3a,3b), which is driven by active tensions at the apical and basal surfaces of the epithelium. We use elements from the theory of active surfaces [1].

#### 1.1 Description of the *Drosophila* embryo as a time-dependent 1D manifold

For simplicity, we describe the *Drosophila* embryo as a time-dependent 1D manifold $\boldsymbol{x}(s,T)$ that follows the epithelial midline in a mid-sagittal section of the embryo. For given position $s$ and time $T$, $\boldsymbol{x}$ is a position in 2D euclidean space, which corresponds to the mid-saggital section when seen from the embryo's right side (as in Fig. 1b,c in the main text). It is parameterized by a scalar $s$ such that the manifold is a loop and runs in clockwise sense with increasing $s$, such that it successively passes through dorsal, anterior, ventral, and posterior part of the embryo, respectively.

Based on the manifold $\boldsymbol{x}(s,T)$, we introduce for given $s$ and $T$ the tangent vector $\boldsymbol{e}$ and length $e$, unit normal vector pointing outside $\boldsymbol{n}$, and local curvature $c$ following standard definitions:

$$\boldsymbol{e} = \partial_s \boldsymbol{x} \tag{S1}$$

$$e = |\boldsymbol{e}| \tag{S2}$$

$$\boldsymbol{n} = \frac{1}{e}\boldsymbol{\varepsilon} \cdot \boldsymbol{e} \tag{S3}$$

$$c = -\frac{1}{e^2}(\partial_s \boldsymbol{e}) \cdot \boldsymbol{n}. \tag{S4}$$

Here, the tensor $\boldsymbol{\varepsilon}$ is the generator of counter-clockwise rotations

$$\boldsymbol{\varepsilon} = \begin{pmatrix} 0 & -1 \\ 1 & 0 \end{pmatrix} \tag{S5}$$

and $\cdot$ denotes the inner product.

Apical and basal surfaces of the embryo are then, respectively:

$$\boldsymbol{x}^{a/b}(s,T) = \boldsymbol{x}(s,T) \pm \frac{h}{2}\boldsymbol{n}(s,T), \tag{S6}$$

where the superscripts $a$ and $b$ correspond to apical and basal surface, and to the signs $+$ and $-$ on the right-hand side, respectively. The variable $h$ denotes the epithelial height. The corresponding tangential vectors are:

$$\boldsymbol{e}^{a/b} = \partial_s \boldsymbol{x}^{a/b} = \left(1 \pm \frac{hc}{2}\right)\boldsymbol{e}. \tag{S7}$$

In the second step, we inserted Eqs. (S6) and used the relation $\partial_s \boldsymbol{n} = c\boldsymbol{e}$. Moreover, we ignored spatial variations in epithelial height $h$ here (Extended Data Fig. 3c).

#### 1.2 Force and torque balance

To define the tension $\boldsymbol{t}$ at some position $s$ of the embryo, we consider an imaginary interface at $s$ that is orthogonal to the manifold $\boldsymbol{x}(s,T)$. Then, $\boldsymbol{t}$ is defined as the force that the part of the embryo behind this interface (larger $s$) exerts on the part of the embryo in front of this interface (smaller $s$). We denote tangential and normal components of $\boldsymbol{t}$ by:

$$t = \frac{1}{e}\boldsymbol{e}\cdot\boldsymbol{t} \tag{S8}$$

$$t_n = \boldsymbol{n}\cdot\boldsymbol{t}. \tag{S9}$$

Analogously, we define the moment $m$ at position $s$ as the torque that the portion of the embryo behind the interface at $s$ exerts on the portion of the embryo in front of the interface. The variable $m$ corresponds thereby to the torque component perpendicular to the mid-sagittal plane (from right to left side of the embryo). We do not consider any other torque component in our 1D model here.

We consider three kinds of external forces that are applied on the embryo: (i) a force density $f^a$ describing friction with the vitelline membrane, which acts tangentially on the apical surface, (ii) a normal force density $-p^a$ acting on the apical surface, which corresponds to the normal force by the vitelline membrane (where the embryo touches the vitelline membrane) or the pressure in the perivitelline space (where the embryo does not touch the vitelline membrane), and (iii) a normal force density $p^b$ that corresponds to the yolk pressure. We ignore here a tangential force on the basal surface by yolk viscosity (see subsection 3.2). Ignoring inertia, force and torque balance in terms of $t$, $t_n$, and $m$ are then (appendix A):

$$t' + ct_n = -\left(1 + \frac{hc}{2}\right) f^a \qquad \text{(tangential force)} \tag{S10}$$

$$t_n' - ct = -\Delta p + \bar{p}hc \qquad \text{(normal force)} \tag{S11}$$

$$m' - t_n = -\frac{h}{2}\left(1 + \frac{hc}{2}\right) f^a, \qquad \text{(torque)} \tag{S12}$$

where the prime denotes the arc-length derivative, $q' := (\partial_s q)/e$ for any $q$, $\Delta p = p^b - p^a$ is the pressure difference across the epithelium and $\bar{p} = (p^a + p^b)/2$ is the average pressure.

To obtain our model equation, Eq. (3) in the main text, force and torque balance need to be complemented by constitutive relations, which link embryonic tensions and moments to deformation, deformation rates, and active apical and basal tensions. To derive these, we follow a virtual work approach. This allows us to properly take active apical an basal tensions into account.

#### 1.3 Virtual work

We consider virtual displacements $\delta\boldsymbol{x}(s,T)$ of the embryo from $\boldsymbol{x}(s,T)$ to $\boldsymbol{x}'(s,T) = \boldsymbol{x}(s,T) + \delta\boldsymbol{x}(s,T)$. These virtual displacements induce virtual mechanical work exerted by active apical and basal tensions $\delta W_{\text{act}}$, work by externally applied forces and torques $\delta W_{\text{ext}}$, a change of an effective bending energy $\delta U_{\text{bend}}$, and dissipated heat $\delta W_{\text{diss}}$. Without inertia, we have:

$$\delta W_{\text{act}} + \delta W_{\text{ext}} = \delta U_{\text{bend}} + \delta W_{\text{diss}}. \tag{S13}$$

We now derive expressions for each of these contributions.

##### 1.3.1 Mechanical work by active apical and basal tensions, $\delta W_{\text{act}}$

The mechanical work by active apical and basal tensions is:

$$\begin{aligned} \delta W_{\text{act}} &= -\oint t^a_{act}\frac{\delta e^a}{e^a}\, e^a \mathrm{d}s - \oint t^b_{act}\frac{\delta e^b}{e^b}\, e^b \mathrm{d}s \\ &= -\oint t^a_{act}\delta e^a \,\mathrm{d}s - \oint t^b_{act}\delta e^b \,\mathrm{d}s. \end{aligned} \tag{S14}$$

Here, $t^a_{act}$ and $t^b_{act}$ are apical and basal active tensions, respectively, $\delta e^a/e^a$ and $\delta e^b/e^b$ are local strain in apical and basal surfaces, and $e^a\mathrm{d}s$ and $e^b\mathrm{d}s$ are infinitesimal apical and basal length elements. From Eqs. (S7) follows:

$$\delta e^{a/b} = \left(1 \pm \frac{hc}{2}\right)\delta e \pm \frac{h}{2} e\delta c. \tag{S15}$$

Insertion into Eq. (S14) yields:

$$\delta W_{\text{act}} = -\oint \left[(t_{act} + cm_{act})\frac{\delta e}{e} + m_{act}\delta c\right] e\mathrm{d}s, \tag{S16}$$

where we introduced (total) active tension $t_{act}$ and active moment $m_{act}$ as:

$$t_{act} = t^a_{act} + t^b_{act} \tag{S17}$$

$$m_{act} = \frac{h}{2}(t^a_{act} - t^b_{act}). \tag{S18}$$

###### 1.3.2 Mechanical work by external forces and torques, $\delta W_{\rm ext}$

The external forces introduced in subsection 1.2 exert the following mechanical work on the embryo:

$$\delta W_{\rm ext} = \oint [f^a \delta x^a_t - p^a \delta x^a_n]\, e^a {\rm d}s + \oint p^b \delta x^b_n\, e^b {\rm d}s, \tag{S19}$$

where $\delta x^a_t := \delta \boldsymbol{x}^a \cdot \boldsymbol{e}/e$ and $\delta x^{a/b}_n := \delta \boldsymbol{x}^{a/b} \cdot \boldsymbol{n}$ are tangential and normal component of the virtual displacements of apical and basal surface, respectively.

Using local force and torque balance, Eqs. (S10)–(S12), this virtual work can also be expressed in terms of tangential tensions $t$ and moments $m$ only (appendix B):

$$\delta W_{\rm ext} = \oint \left[(t + cm)\frac{\delta e}{e} + m\delta c\right] e{\rm d}s. \tag{S20}$$

###### 1.3.3 Effective bending energy, $U_{\rm bend}$

With an effective bending rigidity $\kappa$, the total effective bending energy of the embryo is

$$U_{\rm bend} = \oint \frac{1}{2}\kappa c^2\, e{\rm d}s. \tag{S21}$$

Its variation as a consequence of the virtual displacements $\delta \boldsymbol{x}$ is:

$$\delta U_{\rm bend} = \oint \left[\frac{1}{2}\kappa c^2 \frac{\delta e}{e} + \kappa c \delta c\right] e{\rm d}s. \tag{S22}$$

Here, we assumed that the local bending rigidity $\kappa$ does not change when the tissue is strained.

###### 1.3.4 Dissipated heat, $\delta W_{\rm diss}$

In our model, we assume that dissipation *within the embryo* occurs only due to viscous friction in tangential direction:

$$\delta W_{\rm diss} = \oint \eta u_t \frac{\delta e}{e}\, e{\rm d}s. \tag{S23}$$

Here, $\eta$ is an effective 1D tissue viscosity, and $u_t$ is the tangential strain rate. This tangential strain rate is related to the tangential and normal velocity components $\mathrm{v}$ and $\mathrm{v}_n$ as [1]:

$$u_t = \mathrm{v}' + c\mathrm{v}_n. \tag{S24}$$

Extended Data Fig. 3a,3b shows that the contribution by the normal motion is negligible in our case, so:

$$u_t \simeq \mathrm{v}'. \tag{S25}$$

#### 1.4 Constitutive relations

Inserting all contributions, Eqs. (S16), (S23), (S22) and (S23) with Eq. (S25), into Eq. (S13) and comparing the coefficients in front of $\delta c$, we obtain:

$$m = \kappa c + m_{act}. \tag{S26}$$

Comparing the coefficients in front of $\delta e$, we obtain:

$$t + cm = \frac{1}{2}\kappa c^2 + \eta \mathrm{v}' + t_{act} + cm_{act}. \tag{S27}$$

Inserting (S26), we find:

$$t = \eta \mathrm{v}' + t_{act} - \frac{1}{2}\kappa c^2. \tag{S28}$$

The last term can be interpreted as a tendency of the tissue to leave regions with a high curvature to reduce its bending energy.

#### 1.5 Model equation

To obtain our model equation, we combine tangential force balance, Eq. (S10), with torque balance, Eq. (S12):

$$t' + cm' = -\left(1 + \frac{hc}{2}\right)^2 f^a. \tag{S29}$$

Inserting the constitutive relations for $t$ and $m$, Eqs. (S28) and (S26), we obtain:

$$\eta \mathrm{v}'' + t'_{act} + cm'_{act} = -\left(1 + \frac{hc}{2}\right)^2 f^a. \tag{S30}$$

Here, we assumed homogeneous tissue viscosity $\eta$ and bending rigidity $\kappa$.

Note that using $t_{act}$ and $m_{act}$ from Eqs. (S17) and (S18) just as ad-hoc expressions for active tension and active torque in a formalism such as Ref. [1] can lead to the wrong equation. The deeper reason for this is that the active tension in Ref. [1] corresponds to the virtual work performed by linear strain for constant curvature, which corresponds to $t_{act} + cm_{act}$ (compare Eq. (S16)), while $t_{act}$ is the virtual work performed by linear strain for *zero* curvature.

We set the external force acting tangentially at the apical surface to be a simple substrate friction with the vitelline membrane:

$$f^a = -\gamma \mathrm{v}. \tag{S31}$$

Insertion in Eq. (S30) yields:

$$\eta \mathrm{v}'' + t'_{act} + cm'_{act} = \left(1 + \frac{hc}{2}\right)^2 \gamma \mathrm{v}. \tag{S32}$$

The prefactor in front of the substrate friction comes from two effects that add each the same factor of $(1 + hc/2)$. First, the friction force $f^a$ is a force per length, and it acts on the apical surface, which is by a factor of $(1+hc/2)$ longer than the midline (see also (S10)). Second, since the friction force acts on the apical surface instead of the midline, it locally exerts a torque on the embryo (see also (S12)), which enters the tangential force balance when eliminating the normal tension $t_n$.

In our system, we find that $hc$ is smaller than 1, even though it can reach $\approx$ 0.4 at the poles (Extended Data Fig. 3d). However, at the poles, the epithelium is often further apart from the eggshell, without any noticeable impact on the flow. For simplicity, we thus absorb the factors $(1+hc/2)$ on the right hand side of Eq. (S32) into a homogeneous friction coefficient $\gamma$. Rearranging the terms, we thus have:

$$\eta \mathrm{v}'' - \gamma \mathrm{v} = -t'_{act} - cm'_{act}. \tag{S33}$$

This is equation (3) in the main text (in Fig. 5e). Equation (1) (in Fig. 3a) follows from leaving away the last term on the right-hand side in Eq. (S33), and equation (2) (Fig. 4a) results from including a locally increased friction.

### 2 Emergence of polarized flow

To illustrate the fundamental mechanism driving polarized flow in our model (Fig. 5a-d in the main text), we focus on the simplified situation where the pressure difference $\Delta p$ is large enough to prevent any invagination. In other words, the embryo midline follows a time-independent curve $\boldsymbol{x}_{vm}(s)$ prescribed by the vitelline membrane. If we additionally assume for simplicity that the embryo is incompressible and $s$ is an arc-length coordinate, we have:

$$\boldsymbol{x}(s,T) = \boldsymbol{x}_{vm}\big(s + s_e(T)\big). \tag{S34}$$

In other words, the configuration of the embryo can be entirely described by the time dependence of $s_e(T)$, which describes how the embryo shifts around within the vitelline membrane.

In this case, the interaction between active moment $m_{act}$ and curvature of the vitelline membrane creates an effective force $F_{act}$ that tends to move the whole epithelium in clockwise direction. To see this, we note that such a force corresponds to a virtual work $\delta W_{\mathrm{act}} = F_{act}\delta s_e$. Since the embryo experiences no strain in tangential direction, the virtual mechanical work by apical and basal tensions, Eq. (S16), is:

$$\delta W_{\mathrm{act}} = -\oint m_{act}(s)\delta c(s)\,\mathrm{d}s, \tag{S35}$$

where $e = 1$ since $s$ is arc length variable here. Using $\delta c(s) = c'_{vm}(s + s_e)\delta s_e$, where $c_{vm}$ is the local curvature corresponding to $\boldsymbol{x}_{vm}$, we get:

$$\delta W_{\mathrm{act}} = -\delta s_e \oint m_{act}(s)c'_{vm}(s + s_e)\,\mathrm{d}s. \tag{S36}$$

We thus obtain for $F_{act}$:

$$F_{act} = -\oint m_{act}(s) c'_{vm}(s + s_e)\, \mathrm{d}s. \tag{S37}$$

Or, using a partial integration:

$$F_{act} = \oint m'_{act}(s) c_{vm}(s + s_e)\, \mathrm{d}s. \tag{S38}$$

Note that this corresponds to the integral of the left-hand side of the tangential force balance equation, Eq. (S30).

### 3 Orders of magnitude

#### 3.1 Speed of polarized flow

To compute the speed of the polarized flow, we use the scenario discussed in section 2, i.e. the pressure is large enough for the embryo to be entirely in contact with the embryo, and the embryo is incompressible in tangential direction. Moreover, we consider here a sagittal section with lateral width $\Delta z$, centered around the mid-sagittal plane.

To obtain a rough order of magnitude for the velocity of the polarized flow, we consider an active moment profile of $m_{act}(s) = m^0_{act}$ for $s \in [s_1, s_2]$, and otherwise $m_{act}(s) = 0$. Then we obtain for the effective force $F_{act}$ driving the polarized flow, using Eq. (S38):

$$F_{act} = m^0_{act} \Delta c, \tag{S39}$$

where $\Delta c = c_1 - c_2$ with $c_1 := c(s_1 + s_e)$ and $c_2 := c(s_2 + s_e)$. The active moment results in our system from an active tension $t^0_{act}$ that appears apically, and with Eq. (S18):

$$F_{act} = t^0_{act} \frac{h \Delta c}{2}. \tag{S40}$$

We equate this force with a friction force $F_{ext} = \alpha \Delta z L \bar{\mathrm{v}}$ against the vitelline membrane, where $L$ is the total length of the embryo and $\alpha = \gamma / \Delta z$ is the friction coefficient between embryo and vitelline membrane. We thus obtain for the average tangential speed $\bar{\mathrm{v}}$:

$$\bar{\mathrm{v}} = \frac{t^0_{act}}{\Delta z L \alpha} \frac{h \Delta c}{2}. \tag{S41}$$

With $t^0_{act}/\Delta z \sim 30\,\mathrm{pN} \cdot \mu\mathrm{m}^{-1}$ (tension of myosin-enriched cell-cell interface in the embryo $\sim 300\,\mathrm{pN}$ [2] and cell size $\sim 10\,\mu\mathrm{m}$), $L \approx 10^3\,\mu\mathrm{m}$, and $h\Delta c \sim 0.3$ (Extended Data Fig. 3d), and a friction right after cellularization of $\alpha \approx 3\,\mathrm{pN} \cdot \mathrm{s} \cdot \mu\mathrm{m}^{-3}$ [3], we obtain

$$\bar{\mathrm{v}} \sim 0.1\,\mu\mathrm{m} \cdot \mathrm{min}^{-1}. \tag{S42}$$

This suggests that a curvature-to-active-moment coupling would be sufficient to drive the flow with average speed $\bar{\mathrm{v}} \sim 1\,\mu\mathrm{m}\cdot\mathrm{min}^{-1}$ when the friction with the egg shell decreases by around an order of magnitude, consistent with our quantitative fits.

##### 3.2 Effect of yolk viscosity

In our modeling we have neglected the yolk viscosity, because its effect can be neglected as compared to the friction of the embryo with the vitelline membrane. The friction coefficient between embryo and vitelline membrane right after cellularization has been determined to be $\alpha \approx 2\ldots 3\,\mathrm{pN}\cdot\mathrm{s}\cdot\mu\mathrm{m}^{-3}$ [3].

To compare this to the mechanical effect of the yolk viscosity on the embryo, we consider a situation where a velocity difference of $\Delta\mathrm{v}$ between dorsal and ventral part of the embryo create a simple shear flow with shear rate $\Delta\mathrm{v}/H$ in the yolk, where $H \approx 50\,\mu\mathrm{m}$ is the distance between basal surfaces of dorsal and ventral parts of the embryo. This shear flow leads to a friction force density of $f_Y = \eta_Y \Delta\mathrm{v}/H$, where the yolk viscosity was measured to be $\eta_Y \approx 1\,\mathrm{Pa}\cdot\mathrm{s}$ [4]. The yolk thus exerts a friction force density of maximally $f_Y/\Delta v \approx 0.02\,\mathrm{pN}\cdot\mathrm{s}\cdot\mu\mathrm{m}^{-3}$. This is two orders of magnitude smaller than the friction forces between embryo and vitelline membrane right after cellularization.

#### 4 Model fitting and prediction

##### 4.1 Retrograde flow

To discuss retrograde flow, we first note that equation (2) in Fig. 4a of the main text results from Eq. (S33) by neglecting the last term and spatially modulating friction:

$$\eta \mathrm{v}'' - (1 + g\Theta_G)\gamma \mathrm{v} = -t'_{act}, \tag{S43}$$

where

$$\Theta_G(s) = \begin{cases} 1 & \text{if } s \in G \\ 0 & \text{if } s \notin G \end{cases} \tag{S44}$$

with $G$ being a small region posterior to the apical myosin patch (see Fig. 4a).

Formally, the existence of retrograde flow and its magnitude follows from integrating Eq. (S43) over the whole domain of the embryo:

$$\bar{\mathrm{v}} = -\frac{g\ell_G}{L}\bar{\mathrm{v}}_G. \tag{S45}$$

Here, $\bar{\mathrm{v}}$ is the tangential velocity averaged over the whole epithelium, $L$ is the length of the whole epithelium, $\ell_G$ is the length of region $G$, and $\bar{\mathrm{v}}_G$ is the tangential velocity averaged over region $G$. From Eq. (S45) with $g > 0$ follows directly that overall average flow $\bar{\mathrm{v}}$ and $\bar{\mathrm{v}}_G$ have opposite sign, implying retrograde flow within $G$.This is because frictional force in the high friction ($g > 0$)

region must be balanced by frictional force in the other, low friction ($g = 0$) region. Since frictional force is proportional to velocity and total force must sum to zero, this means that if the velocity in the low friction region is positive (clockwise), the velocity in the region of high friction will have to be negative (counterclockwise). Moreover, the absence of localized friction, $g = 0$, Eq. (S45) implies zero average velocity, $\bar{\mathrm{v}} = 0$.

#### 4.2 Model Fitting

To quantitatively compare equations (1)–(3) in the main text (which follow from Eq. (S33)) to experimental data, we assumed a linear relation between apical and basal active tension, $t^a_{act}$ and $t^b_{act}$, and the respective sqh::GFP signal, $I_a$ and $I_b$:

$$t^a_{act} = f_a I_a \qquad\qquad t^b_{act} = f_b I_b. \tag{S46}$$

We assume that $f_a$ and $f_b$ can be different, where due to the different cytoskeletal structures apically and basally, we expect $f_a > f_b$. According to Eqs. (S17) and (S18), we thus have:

$$t_{act} = f_a I_a + f_b I_b \tag{S47}$$

$$m_{act} = \left(f_a I_a - f_b I_b\right)\frac{h}{2}. \tag{S48}$$

Insertion into equation (3) in the main text (i.e. Eq. (S33)) and division by $\eta$ yields:

$$\mathrm{v}'' - \frac{1}{l_H^2}\mathrm{v} = -r_a I_a' \left(1 + \frac{ch}{2}\right) - r_b I_b' \left(1 - \frac{ch}{2}\right), \tag{S49}$$

where $l_H = \sqrt{\eta/\gamma}$ is the hydrodynamic length scale, $r_a = f_a/\eta$, and $r_b = f_b/\eta$. The parameters $r_a$ and $r_b$ have units of rates per pixel intensity – they indicate how fast the epithelium contracts per pixel intensity of myosin.

We use Eq. (S49) to fit model equation (3), while using the correspondingly modified equations to fit equations (1) and (2) in the main text. In particular, we leave out the terms $\sim hc$ on the right-hand side for equations (1) and (2), and we introduce a localized friction for equation (2).

To fit Eq. (S49) to experimental data, we first use the measured $c$, $I_a$,$I_b$ and $h$ to numerically solve Eq. (S49) for v at each time step. For this, we discretize this equation in space with regular lattice spacing $\Delta s = 0.01$ and solve the resulting linear equation in $\mathrm{v}(s)$ in python using a sparse matrix inverter.

To obtain the parameters $l_H$, $r_a$, and $r_b$ (and $g$ for equation (2)), we always fitted the the theoretical predictions for $\mathrm{v}(s)$ to its respective measured curves $\mathrm{v}(s)$ *at all time points simultaneously* between $T_{asb} = -5 \ldots 8$ min. To this end, we used the python routine curve-fit to minimize the squared distances between theoretical and measured v summed over all positions and times (minimal $\chi^2$ plotted in Fig. 5i). In particular, we carried out two kinds of simultaneous fits. First, for many fits, we imposed that all parameter values should be the same at all time points. These corresponds to the blue curves in Fig. 3c,e, 4d,e, and 5g,h. Second, for some fits, we allowed $l_H$ to be different for each time point, while we imposed that all other parameters have to be the same value at each time point. This corresponds to the magenta curves in Fig. 5g,h and fit curves in Extended Data Fig. 6,7,8,10.

#### 4.3 Limitation on compression of apical myosin patch

We realized that our fitting of model equation (3) resulted in a substantial increase of the hydrodynamic length scale of over an order of magnitude around the time of symmetry breaking (Extended Data Fig. 6d). Upon close examination of possible causes for this jump, we first noted that assuming a dominant role of apical myosin during the asymmetric phase, the velocity of polarized flow should scale as $\bar{\mathrm{v}} \sim f_a I_a/\gamma = I_a r_a l_H^2$. Second, we noticed a steep decrease in velocity in our velocity fit curves in the region where the apical myosin patch is, around $s \approx 0.03$ (magenta curves in Fig. 5i, blue region in Extended Data Fig. 6e). This velocity decrease in the fit curves corresponds to a strong contractile flow, which is created by apical myosin and resisted by tissue viscosity. The corresponding contraction rate is of order $\sim f_a I_a/\eta = I_a r_a$. Taken together, in our $\chi^2$ minimization based fitting, to keep the contraction of the apical myosin patch close to measured values while keeping large enough $\bar{\mathrm{v}}$, the hydrodynamic length scale $l_H$ needs to be large during the asymmetric phase.

To test these ideas, we also examined a model where the contraction rate of the region with the apical myosin patch (the primordium) would be limited. Limiting this contraction rate makes sense, because the primordium undergoes isotropic contraction. Indeed, this region around the apical myosin distribution has increased epithelial height in the asymmetric phase (Extended Data Fig. 6f). There will thus be a limit on how far this part of the tissue will be able to contract until elastic resistance prevents further contraction. In our model, we do not have included elasticity, which would require including an additional parameter and defining reference states. To circumvent these issues and keep the model simple, we have decided to study the consequences of a limited promordium contraction rate in a symmetric region around the peak of apical myosin distribution at $s = 0.03$ (Extended Data Fig. 3f) by substantially increasing viscosity in the primordium region (Eq. 4 in Extended Data Fig. 7a).

We show results of fits where we locally increased viscosity by a factor of $e = 100^{**}$ with varying length $0.08 \geq L_E \geq 0.0$, i.e restricting within the apical myosin domain. We find that the contraction rate in the primordium region is indeed decreased (Extended Data Fig. 7b). Moreover, the corresponding increase in hydrodynamic length scale is also much smaller now, from $l_H \approx 0.04$ (40 $\mu m$) during symmetric flow to $l_H \approx 0.4$ (400 $\mu m$)during asymmetric flow (Extended Data Fig. 7c). Taken together, taking into account a limited primordium contraction rate, our data can be explained better (smaller $\chi^2$, see Extended Data Fig. 7d) and with a smaller decrease in friction with the eggshell.

**We find that our results are largely independent of the increase in viscosity as long as $e > 10$ (Extended Data Fig. 8).

##### 4.4 Simulations using a simplified model

To obtain a better intuition, we simplified the embryo by representing its shape as an ellipse (elliptic contour in Extended Data Fig. 5a,6a), discretized by 100 evenly spaced nodes. For these simulations we neglected basal myosin and approximated the distribution of apical myosin by a rectangular function (green patch in Extended Data Fig. 5b,6b) with height $I_a^{sim}$. To simulate equation (2), we moreover approximated the patch of increased friction, $\Theta_G$, by another rectangular function (magenta patch in Extended Data Fig. 5b) that advected with the epithelium. In all simulations, we choose $I_a^{sim}$ to be of the order of experimentally measured apical myosin intensity $I_a$ in Fig. 3b,d, and the values of the physical parameters were chosen from the fit values to the experimental data (see Table, Extended Data Fig. 9x).

We then simulated discrete time steps $\Delta t = 0.5\,\text{min}$, where at a given time point $T_{sim}$, we solved equation (2) or (3) for velocity using the python solver described in the previous section to obtain the velocity field $\text{v}_{sim}(s)$. To further simplify our simulation, we did not allow for any deformation of the epithelium. We thus advanced the whole epithelium at each time step by the distance $\bar{\text{v}}\Delta t$. This introduced a time dependence in our solution for equation (3), due to a changing offset between curvature $c(s)$ and myosin profile $I_a^{sim}(s)$.

### Appendix

#### A Force and torque balance

To derive the force and torque balance relations, Eqs. (S10)–(S12), we roughly follow the approach from [1].

We start from noting that in an overdamped system, the total force acting on any piece of the embryo between $s_1$ and $s_2$ needs to vanish:

$$\begin{aligned} 0 = \boldsymbol{t}(s_2) - \boldsymbol{t}(s_1) + \int_{s_1}^{s_2} \left[ f^a \frac{1}{e^a} \boldsymbol{e}^a - p^a \boldsymbol{n}^a \right] e^a \mathrm{d}s \\ + \int_{s_1}^{s_2} p^b \boldsymbol{n}^b \, e^b \mathrm{d}s \end{aligned} \tag{A50}$$

Here, the terms on the right hand side are the force exerted by the region of the part of embryo behind $s_2$, the force exerted by the part of the embryo before $s_1$, the external force exerted on the apical surface, and the external force exerted on the basal surface.

The derivative of Eq. (A50) with respect to $s \equiv s_2$ is:

$$0 = \partial_s \boldsymbol{t} + f^a \boldsymbol{e}^a + (p^b e^b - p^a e^a) \boldsymbol{n} \tag{A51}$$

Using Eqs. (S7) and the arc-length derivative, $q' := (\partial_s q)/e$ for any $q$:

$$0 = \boldsymbol{t}' + f^a \left(1 + \frac{hc}{2}\right) \frac{1}{e}\boldsymbol{e} + (\Delta p - \bar{p}hc)\boldsymbol{n}. \tag{A52}$$

Here, we have defined $\Delta p = p^b - p^a$ and $\bar{p} = (p^a + p^b)/2$. Using $\boldsymbol{t} = t\boldsymbol{e}/e + t_n\boldsymbol{n}$ together with the relations $\boldsymbol{n}' = c\boldsymbol{e}/e$ and $(\boldsymbol{e}/e)' = -c\boldsymbol{n}$, Eq. (A52) becomes:

$$\begin{aligned} 0 = {}& t'\frac{1}{e}\boldsymbol{e} - ct\boldsymbol{n} + t_n'\boldsymbol{n} + ct_n\frac{1}{e}\boldsymbol{e} \\ &+ f^a \left(1 + \frac{hc}{2}\right)\frac{1}{e}\boldsymbol{e} + (\Delta p - \bar{p}hc)\boldsymbol{n}. \end{aligned} \tag{A53}$$

Tangential and normal force balance, Eqs. (S10) and Eq. (S11), can now be directly read off directly from tangential and normal part of this equation.

The total torque acting on the same piece of embryo also needs to vanish:

$$\begin{aligned} 0 = {}& m(s_2) + \boldsymbol{x}(s_2)\cdot\boldsymbol{\varepsilon}\cdot\boldsymbol{t}(s_2) \\ &- m(s_1) - \boldsymbol{x}(s_1)\cdot\boldsymbol{\varepsilon}\cdot\boldsymbol{t}(s_1) \\ &+ \int_{s_1}^{s_2} \boldsymbol{x}^a\cdot\boldsymbol{\varepsilon}\cdot\left[f^a\frac{1}{e^a}\boldsymbol{e}^a - p^a\boldsymbol{n}^a\right] e^a \mathrm{d}s \\ &+ \int_{s_1}^{s_2} \boldsymbol{x}^b\cdot\boldsymbol{\varepsilon}\cdot p^b\boldsymbol{n}^b\, e^b \mathrm{d}s \end{aligned} \tag{A54}$$

The derivative with respect to $s \equiv s_2$ is:

$$\begin{aligned} 0 = {}& \partial_s m + \boldsymbol{e}\cdot\boldsymbol{\varepsilon}\cdot\boldsymbol{t} + \boldsymbol{x}\cdot\boldsymbol{\varepsilon}\cdot(\partial_s\boldsymbol{t}) \\ &+ \boldsymbol{x}^a\cdot\boldsymbol{\varepsilon}\cdot[f^a\boldsymbol{e} - p^a e^a\boldsymbol{n}] \\ &+ \boldsymbol{x}^b\cdot\boldsymbol{\varepsilon}\cdot p^b e^b\boldsymbol{n}. \end{aligned} \tag{A55}$$

After using Eqs. (S3) and (S9) as well as consecutive insertion of Eqs. (S6) and (A51):

$$0 = \partial_s m - e t_n + \frac{h}{2} f^a \boldsymbol{n}\cdot\boldsymbol{\varepsilon}\cdot\boldsymbol{e}^a \tag{A56}$$

Insertion of Eq. (S7) yields torque balance, Eq. (S12):

$$0 = m' - t_n + \frac{h}{2}\left(1 + \frac{hc}{2}\right) f^a. \tag{A57}$$

### B Virtual work by external forces and torques

Using force and torque balance, we show here that the expressions in Eqs. (S19) and (S20) for the virtual work by external forces and torques are equivalent [1]. To this end, we start with expression Eq. (S20):

$$\delta W_{\mathrm{ext}} = \oint \left[(t + cm)\frac{\delta e}{e} + m\delta c\right] e\mathrm{d}s. \tag{B58}$$

Using

$$\delta e = \frac{1}{e}\boldsymbol{e}\cdot\partial_s\delta\boldsymbol{x} \tag{B59}$$

$$\delta c = -\frac{2c}{e^2}\boldsymbol{e}\cdot\partial_s\delta\boldsymbol{x} - \frac{1}{e}\boldsymbol{n}\cdot\partial_s\left(\frac{1}{e}\partial_s\delta\boldsymbol{x}\right), \tag{B60}$$

Eq. (B58) becomes:

$$\begin{aligned}\delta W_{\text{ext}} = &\oint (t - cm)\frac{1}{e}\boldsymbol{e}\cdot\partial_s\delta\boldsymbol{x}\,\mathrm{d}s \\ &- \oint m\boldsymbol{n}\cdot\partial_s\left(\frac{1}{e}\partial_s\delta\boldsymbol{x}\right)\mathrm{d}s.\end{aligned} \tag{B61}$$

After partial integrations:

$$\begin{aligned}\delta W_{\text{ext}} = &-\oint\left(\partial_s\left[(t - cm)\frac{1}{e}\boldsymbol{e}\right]\right)\cdot\delta\boldsymbol{x}\,\mathrm{d}s \\ &- \oint\left(\partial_s\left[\frac{1}{e}\partial_s[m\boldsymbol{n}]\right]\right)\cdot\delta\boldsymbol{x}\,\mathrm{d}s.\end{aligned} \tag{B62}$$

Using $\boldsymbol{n}' = c\boldsymbol{e}/e$:

$$\delta W_{\text{ext}} = -\oint\left(t\frac{1}{e}\boldsymbol{e} + m'\boldsymbol{n}\right)'\cdot\delta\boldsymbol{x}\,e\mathrm{d}s. \tag{B63}$$

Using both $\boldsymbol{n}' = c\boldsymbol{e}/e$ and $(\boldsymbol{e}/e)' = -c\boldsymbol{n}$:

$$\delta W_{\text{ext}} = -\oint\left(t'\frac{1}{e}\boldsymbol{e} - ct\boldsymbol{n} + m''\boldsymbol{n} + cm'\frac{1}{e}\boldsymbol{e}\right)\cdot\delta\boldsymbol{x}\,e\mathrm{d}s. \tag{B64}$$

Combining tangential and normal force balance respectively with torque balance, Eqs. (S10)–(S12), we have:

$$t' + cm' = -\left(1 + \frac{hc}{2}\right)^2 f^a \tag{B65}$$

$$m'' - ct = -\left[\frac{h}{2}\left(1 + \frac{hc}{2}\right)f^a\right]' - \Delta p + \bar{p}hc. \tag{B66}$$

Insertion in (B64) yields:

$$\begin{aligned}\delta W_{\text{ext}} = &\oint\left(1 + \frac{hc}{2}\right)^2 f^a\frac{1}{e}\boldsymbol{e}\cdot\delta\boldsymbol{x}\,e\mathrm{d}s \\ &+ \oint\left(\partial_s\left[\frac{h}{2}\left(1 + \frac{hc}{2}\right)f^a\right]\right)\boldsymbol{n}\cdot\delta\boldsymbol{x}\,\mathrm{d}s \\ &+ \oint(\Delta p - \bar{p}hc)\boldsymbol{n}\cdot\delta\boldsymbol{x}\,e\mathrm{d}s.\end{aligned} \tag{B67}$$

After partial integration of the second integral:

$$\begin{aligned}
\delta W_{\mathrm{ext}} = & \oint \left(1+\frac{hc}{2}\right)^2 f^a \frac{1}{e}\boldsymbol{e}\cdot\delta\boldsymbol{x}\, e\mathrm{d}s \\
& - \oint \frac{h}{2}\left(1+\frac{hc}{2}\right) f^a (\boldsymbol{n}\cdot\delta\boldsymbol{x})'\, e\mathrm{d}s \\
& + \oint \left(1-\frac{hc}{2}\right) p^b \boldsymbol{n}\cdot\delta\boldsymbol{x}\, e\mathrm{d}s \\
& - \oint \left(1+\frac{hc}{2}\right) p^a \boldsymbol{n}\cdot\delta\boldsymbol{x}\, e\mathrm{d}s.
\end{aligned} \tag{B68}$$

Using $(\boldsymbol{n}\cdot\delta\boldsymbol{x})' = c(\boldsymbol{e}/e)\cdot\delta\boldsymbol{x} + \boldsymbol{n}\cdot\delta\boldsymbol{x}'$ and $(\boldsymbol{e}/e)\cdot\delta\boldsymbol{n} = -\boldsymbol{n}\cdot\delta\boldsymbol{x}'$, as well as $\boldsymbol{n}\cdot\delta\boldsymbol{n} = 0$, this becomes:

$$\begin{aligned}
\delta W_{\mathrm{ext}} = & \oint \left(1+\frac{hc}{2}\right) f^a \frac{1}{e}\boldsymbol{e}\cdot\left(\delta\boldsymbol{x}+\frac{h}{2}\delta\boldsymbol{n}\right) e\mathrm{d}s \\
& + \oint \left(1-\frac{hc}{2}\right) p^b \boldsymbol{n}\cdot\left(\delta\boldsymbol{x}-\frac{h}{2}\delta\boldsymbol{n}\right) e\mathrm{d}s \\
& - \oint \left(1+\frac{hc}{2}\right) p^a \boldsymbol{n}\cdot\left(\delta\boldsymbol{x}+\frac{h}{2}\delta\boldsymbol{n}\right) e\mathrm{d}s.
\end{aligned} \tag{B69}$$

Using Eqs. (S6) and (S7), as well as $\boldsymbol{n}\cdot\delta\boldsymbol{n} = 0$:

$$\begin{aligned}
\delta W_{\mathrm{ext}} = & \oint f^a \frac{1}{e^a}\boldsymbol{e}^a\cdot\delta\boldsymbol{x}^a\, e^a\mathrm{d}s \\
& + \oint p^b \boldsymbol{n}\cdot\delta\boldsymbol{x}^b\, e^b\mathrm{d}s \\
& - \oint p^a \boldsymbol{n}\cdot\delta\boldsymbol{x}^a\, e^a\mathrm{d}s.
\end{aligned} \tag{B70}$$

This is the same expression as in Eq. (S19).
